# *Echinococcus granulosus* antigen B acts as an LPS-scavenging lipoprotein *in vitro* preventing TLR4-mediated activation of dendritic cells

**DOI:** 10.1101/2025.07.02.662828

**Authors:** Lagos Magallanes S, Beasley Lomazzi A, F Zamarreño, F Carrión, M Fló, J Dutto, J Julve, M Costabel, M Maccioni, AM Folle, AM Ferreira

**Author notes:** Address correspondence to Ana M. Ferreira, and Maite Folle.

## Abstract

*Echinococcus granulosus* sensu lato antigen B (EgAgB) is a major parasite lipoprotein, produced by the hydatid and released at the host-parasite interface. Accumulating evidence supports that EgAgB may exert immunomodulatory effects on myeloid cells; however, the underlying molecular mechanisms remain poorly understood. We examined the impact of native EgAgB (nEgAgB) and recombinant EgAgB8/1 (rEgAgB) on lipopolysaccharide (LPS)-induced activation of bone marrow-derived dendritic cells (BMDC), to help elucidate these mechanisms. Both immunoaffinity-purified nEgAgB or rEgAgB induced modest BMDC activation, indicated by the production of IL-6, IL-12p40, and nitric oxide, but not IFN-β. This activation was primarily attributed to LPS traces in EgAgB preparations since it was nearly abolished by a specific TLR4 inhibitor and in *Tlr4*^-/-^ BMDC, while EgAgB binding to BMDC was TLR4-independent. Notably, both nEgAgB and rEgAgB inhibited LPS-induced cytokine and nitric oxide production, and disrupted TLR4 dimerization and endocytosis. Competitive binding assays showed that EgAgB and human high-density lipoprotein (hHDL) similarly inhibited LPS binding to macrophages and BMDC; however, EgAgB more effectively suppressed LPS-induced cytokine secretion. Contrastingly, EgAgB did not modulate BMDC responses to lipoteichoic acid, unlike hHDL. Using dynamic light scattering and an ELISA-like assay, we demonstrated a higher potential of EgAgB to bind LPS than hHDL. Additionally, docking analyses suggest the presence of a defined LPS-binding interface in EgAgB8/1 subunit. Overall, these findings reveal a novel binding property of EgAgB, which enables it to act as an extracellular LPS scavenger, interfering with TLR4-mediated LPS recognition and downstream proinflammatory responses in myeloid cells.

## Background

*Echinococcus granulosus* sensu lato comprises a complex of cestode species whose larval stages cause cystic echinococcosis (CE), a chronic disease primarily affecting domestic ungulates, with humans serving as accidental hosts. The larva (hydatid) is a highly antigenic structure filled with fluid (HF), which typically develops within host visceral organs, mainly the liver and lungs. To resist host defences, the hydatid effectively modulates the host immune response, likely employing multiple evasion strategies to mitigate inflammation and potentially harmful effector mechanisms. When these strategies are successful, host inflammation is resolved, leading to the formation of a host-derived fibrous layer that surrounds the hydatid and contributes to keeping the parasite away from immune cells (1, 2). In this regard, antigen B (EgAgB) has garnered significant attention for its capacity to modulate host inflammation (reviewed in (3, 4)), although the molecular basis underlying this modulation remains unknown.

EgAgB is a lipoprotein abundantly produced by the hydatid and released into the host-parasite interface as evidenced by its detection in host infected tissues and lymphoid organs by immunohistochemistry (5). The interaction of EgAgB with immune cells is further supported by the presence of specific antibodies in infected patients; in fact, its detection is regarded as a valuable tool for the diagnosis of human CE (6–8). Initially, immunomodulatory studies demonstrated that *in vitro* EgAgB limited the inflammatory response of neutrophils, dendritic cells (DC), and monocyte/macrophages using various agonists for cell activation, including pathogen-associated molecular patterns (PAMPs) such as crude bacterial components, formyl peptides, and lipopolysaccharides (LPS), as well as endogenous and synthetic activators (C5a, platelet-activating factor, phorbol ester) (9–13). Recent studies have extended these modulatory properties to *in vivo* inflammatory models including LPS-induced peritonitis (14), bowel disease (15), sepsis (16) and immune thrombocytopenia (17). Additionally, it has been proposed that EgAgB facilitates the development of M2-type macrophages (F4/80^+^ CD206^+^) (15–17) and a Th2-type immune response, indicated by the cytokine profile (IL-4/IFN-γ ratio) induced by exposing peripheral blood mononuclear cells (PBMC) from CE patients or human DC to EgAgB (10, 11).

During the last few years our research group has been interested in elucidating the molecular and cellular mechanisms underlying the immunomodulatory properties of EgAgB, as these insights may reveal novel molecular targets for controlling inflammation in other inflammatory disorders. With this perspective, we first conducted a comprehensive biochemical characterisation of the composition of native EgAgB (nEgAgB), revealing a high heterogeneity of the lipids physiologically bound to the protein component (including fatty acids, sterols, sterol esters, triacylglycerols, and phospholipids) (18). Regarding EgAgB’s protein moiety, it is encoded by a multigenic and polymorphic family whose orthologs belong to the hydrophobic ligand-binding protein (HLBP) family specific to cestodes. In *E. granulosus* s.s., there are five clades (*EgAgB1-EgAgB5*) that encode ∼8 kDa mature products, referred to as EgAgB8/1 to EgAgB8/5 subunits, which have the ability to oligomerize. We found that EgAgB8/1 was the predominant subunit in nEgAgB purified from the HF of various *E. granulosus* s.l. species, including those belonging to *E. granulosus* s.s. (19). Based on size and chemical composition (protein and lipid), we postulate that nEgAgB assembles as a plasma lipoprotein particle, by concentrating hydrophobic lipids in the core, surrounded by a layer of phospholipids into which protein subunits (∼12-14 subunits per particle) are inserted. Among vertebrate plasma lipoproteins, nEgAgB would be more similar to the human high-density lipoprotein (hHDL, (20)), particularly to its small subfraction (hHDL_3_).

Studies on nEgAgB immunomodulation properties are challenging; although it is abundant among parasite HF components, purifying it from HF in sufficient amounts and quality for these studies has proven difficult (14). Some *in vitro* (10, 11) and *in vivo* (15, 17) studies employed an EgAgB preparation obtained by precipitation, including a boiling treatment that likely denatured the lipoprotein. Despite this, the heat-treated EgAgB shared modulatory effects on LPS-mediated myeloid cell activation with nEgAgB. We recently described the preparation and characterization of the recombinant form of EgAgB8/1 (rEgAgB), which was expressed in insect cells as an alternative for immunological studies. This recombinant form shows a high degree of similarity to nEgAgB in terms of its ability to form a lipoprotein particle of similar size and lipid composition, and its capacity to modulate monocyte and macrophage activation by LPS (14). In this work, we focused on understanding the immunomodulatory properties of the native form of EgAgB on myeloid cells, using DC as a cellular model for their importance in the development of the innate and adaptive immune responses. Interestingly, we found that both nEgAgB and rEgAgB preparations showed the ability to activate bone marrow DC (BMDC) on their own, although they inhibited the LPS-induced activation of BMDC, particularly in terms of cytokine secretion. Our results suggest that these contrasting effects may be due to EgAgB’s ability to bind LPS in the extracellular environment interfering with TLR4-driven cell signalling. This is a binding property shared with human high-density lipoprotein (hHDL) and its smaller fraction hHDL_3_, which have been described as LPS scavengers (21), being hHDL_3_ involved in the protection of the liver from inflammation and fibrosis in mice (22). The biological implications of these findings for CE immunobiology are discussed.

## Methods

### EgAgB preparations

*Native EgAgB (nEgAgB):* nEgAgB was obtained from HF from fertile hydatids of naturally infected cows collected during the routine work of local abattoirs (Montevideo, Uruguay). HF was aspired under aseptic conditions and preserved at -20°C with EDTA 5 mM, 3,5-di-tert-butyl-4-hydroxytoluene (BHT) 20 μM. Protoscoleces were recovered for hydatids genotyping by amplifying and sequencing a fragment of the mitochondrial cytochrome c oxidase subunit 1 (COX1) (23). Hydatids belonged mainly to *E. granulosus* s.s. (G1 genotype) but also to *E. ortleppi* (G5 genotype). To obtain enough nEgAgB in each purification HF was pooled, corresponding to a G1/G5 ratio around 80/20 in volume. nEgAgB was purified by a three-step fractionation protocol based on an ion exchange chromatography on Q-Sepharose, followed by sequential ultracentrifugation in a KBr gradient, and a final immunoaffinity chromatography step using an anti-EgAgB8/1 nanobody (clone 1) coupled to Sepharose (11). All solutions and buffers used during the purification protocol were prepared with pyrogen-free water (ICU-VITA, Uruguay) and the last steps -immunoaffinity followed by desalting on a PD-10 column (GE Healthcare, USA)-were carried out in a laminar flow cabinet. Due to the amount of parasite material (HF) required for including the ultracentrifugation step, for some studies (as indicated in the results) we purified nEgAgB without ultracentrifugation. This alternative approach did not significantly alter the particle size, the SDS-PAGE protein profile, or the immunomodulatory effects observed on myeloid cells (see Supplementary Fig. 1). Moreover, excluding the ultracentrifugation step enabled the hole purification protocol in a sterile environment (laminar flow cabinet).

*Recombinant EgAgB8/1 (rEgAgB):* rEgAgB8/1 (accession number U6JQF4) was produced following the previously optimized protocol (14). Briefly, EgAgB8/1-transfected S2 cells were defrosted and cultured in Xpress medium (Lonza, Switzerland) containing penicillin/streptomycin and puromycin for transfected-cell selection. Cells were grown in a culture flask until a 7×10^6^ concentration was reached and rEgAgB production was induced with CdCl_2_ 5 μM for 7 days. Culture supernatants were recovered and, after pH adjusted to 8 and an overnight incubation at 4°C, were clarified by centrifugation and filtered through 0.22 μm. rEgAgB was purified from culture supernatants on a Strep-Tactin XT-agarose column (IBA, Life Science) following an immunoaffinity chromatography with anti-EgAgB8/1-Sepharose column (11).

EgAgB preparations were finally equilibrated in phosphate buffered saline pH 7.2 (PBS) containing EDTA 5 mM, BHT 20 μM, antibiotic (Capricorn), to prevent further oxidation and contamination, and sucrose 10%, to preserve the lipoprotein structure, (20) (PBS_EBAb_) and were conserved at -80°C until use. All EgAgB preparations were examined in terms of size homogeneity (average hydrodynamic diameter, D_H_) by Dynamic Light Scattering (DLS). To that end, samples (70 μL, 1 mg/mL) were placed in disposable cuvettes (UVette, Eppendorf) and pre-incubated at 25°C before measurement. Data from triplicate measurements were averaged and analysed with Zetasizer Software v7.13 (Malvern Panalytical) to obtain size distribution of the samples (weighted by volume as assuming sphericity and homogeneity of particles). Reported D_H_ values were calculated as the mean value for the main peak. In addition, EgAgB protein component was quantified using the bicinchoninic acid assay (BCA, Thermo) and analysed by SDS-PAGE and Western blot as previously described (14). Endotoxin level was also controlled using a chromogenic *Limulus amoebocyte* lysate assay with a cut-off of 0.03 EU/mL (LAL, Beltrán Zunino, Montevideo). Four and three preparations of nEgAgB and rEgAgB were used along this investigation.

### Cell line and generation of bone marrow derived dendritic cells

The human monocyte-like cell line THP-1 (American Type Culture Collection, ATCC, USA) was cultured in RPMI 1640 medium containing HEPES 10 mM, sodium bicarbonate 1.5 g/L, sodium pyruvate 1 mM, glutamine 2 mM, AbAm and heat inactivated Foetal Calf Serum (FCS, Probiomont, Montevideo, Uruguay) 10% (v/v) at 37 °C in a humidified atmosphere with 5% CO_2_ (v/v), following ATCC recommendations. For macrophage differentiation, cells were stimulated with PMA (50 ng/mL, SIGMA) for 72 h. Murine bone-marrow derived dendritic cells (BMDC) were generated by differentiation of bone marrow precursors following a procedure approved by Comisión Honoraria de Experimentación Animal (CHEA Uruguay, protocol N° 889, Exp 101900-000875-19.1), which was based on a standard method (24). Ten weeks old female from C57BL/6 mice (Institut Pasteur, Montevideo) and from *Tlr4*^-/-^ and *Tlr2*^-/-^ mice (strains #007227 and #004650, respectively, both on C57BL/6 background, Jackson Laboratories) were used. Briefly, on day 0, the bone marrow from tibias and femurs was disaggregated by passage through a 24G syringe. Bone marrow precursors were counted and seeded at 3 x 10^6^ per 100 mm petri dishes in 10 mL of complete medium (RPMI 1640 with FCS 10%, L-glutamine 2 mM, penicillin/streptomycin 100 U/mL and J558 cell line supernatant containing GM-CSF 1% (v/v). On day 3, 10 mL of complete medium was added. On days 5 and 7, 10 ml of the culture medium was removed and added 10 ml of complete medium. On day 10, non-adherent cells were collected, between 85 and 95% being CD11c^+^, and used for stimulation.

### Effects of EgAgB BMDC activation

BMDC derived from C57BL/6 mice were seeded at a density of 0.4 x 10^6^ cells per well (96-well plate) and exposed to different concentrations (1-10 µg/mL) of nEgAgB, rEgAgB, or vehicle (Veh, PBS_EBAb_, as a control) in the presence or absence of LPS 10 ng/mL (*Eschericchia coli* O127:B8, Sigma-Aldrich, #L4516) and incubated at 37 °C in a humidified atmosphere with 5% CO_2_ (v/v). After 18 hours, the culture supernatant was collected to determine the levels of inflammatory cytokines (IL-6, IL-12p40, IFN-β) and nitric oxide (·NO) generation, and the cells were recovered to measure surface receptors by flow cytometry as described below. In additional assays, we compared the modulation capacity of EgAgB and hHDL. To that end, hHDL fraction was isolated from pooled plasma by sequential flotation ultracentrifugation as previously described (25). The obtained hHDL fraction presented a D_H_ 11.2 ± 4.1 nm and an LPS content lower than 0.001 ng/µg protein. BMDC were exposed to a range of concentrations of either nEgAgB (0.5-10 µg/mL) or hHDL (10-100 µg/mL) in the presence or absence of LPS (10 ng/mL) or LTA (0.5 µg/mL, *Staphylococcus aureus,* Invivogen #tlrl-pslta) for 18 hours. BMDC responses were examined by measuring IL-6 and IL-12p40 concentrations in culture supernatants.

### Studies on LPS and TLR4 involvement in EgAgB effects on BMDC activation

To inhibit cell activation induced by LPS traces in samples, nEgAgB or rEgAgB (1-10 µg/mL) were pretreated with polymyxin B (10 µg/mL, Sigma-Aldrich) or cultured medium (control) for 2 hs at 37°C and then used to stimulate BMDC at the conditions described above. In parallel, LPS (10 ng/mL) and Pam3CSK4 (60 ng/mL, Invivogen #tlrl-pms) were analysed as controls. To inhibit TLR4-mediated responses, BMDC were preincubated with the TLR4-specific inhibitor TAK-242 (10 µM, USBiological) for 1 h and then stimulated with nEgAgB/rEgAgB (0.5-10 µg/mL), LPS (10 ng/mL) or Pam3CSK4 (60 ng/mL, as control) for 18 hs. Additionally, EgAgB effects were examined on BMDC derived from wild type (WT), Tlr*4*^-/-^ and *Tlr2*^-/-^mice for comparison. In these studies, EgAgB preparations (0.5-10 µg/mL) were assayed and cell responses examined by quantification of IL-6 and IL-12p40and in culture supernatants.

### Measurement of cell responses

Mouse and human IL-6, IL-12p40 and IFN-β were determined in culture supernatants by a capture ELISA using paired antibodies from BD, Biolegend or R&D. ·NO generation was measured based on their conversion in nitrite (NO ^-^), which was quantified by the colorimetric Griess assay (26). Briefly, cell culture supernatants were transferred (50 μL/well) to 96-well flat-bottom plates and 50 μL of sulphanilamide (Sigma, 1% wt/vol in 2.5% H_3_PO_4_) and 50 μL of naphthyl ethylenediamine dihydrochloride (Sigma, 0.1% wt/vol in 2.5% H_3_PO_4_) were added. After 5 min incubation, the absorbance at 540 nm (A_540_) was measured and converted to nitrite concentration based on a NaNO_2_ standard curve. Surface molecules including CD86, CD40 and CD11c were quantified by flow cytometry. To that end, cells were stained using conventional protocols with Live/DEAD-Green (Invitrogen) and the corresponding fluorescent conjugates diluted in PBS containing BSA 0.1% w/v, EDTA 2 mM (FACS). The appropriate controls -fluorescence minus one (FMO)-were also prepared. The information about all fluorescent conjugates used for flow cytometry is detailed in the Supplementary Table 1. Data were acquired on a FACSCanto II cytometer, and analysed using the FlowJoTM package (Version 7.6.2), gating on Live/DEAD-Green^-^ CD11c^+^ events (gating strategy is shown in Supplementary Fig. 2).

### Studies on TLR4 dependence of EgAgB binding to BMDC

EgAgB binding to BMDC derived from WT, *Tlr4*^-/-^ and *Tlr2*^-/-^ mice was examined by flow cytometry. All incubations and washing steps were carried out in a binding buffer (PBS pH 7.2 containing FCS 1% (v/v), NaN_3_ 0.1% (w/v) and EDTA 2 mM). Cells (0.2 x 10^6^ cells/well in a 96-well V-bottom plate) were incubated for 1 h at 4 °C with increasing concentrations of nEgAgB (1-100 μg/mL) or vehicle (Veh) as a control. After two washing steps, Fc receptors were blocked by the addition of 10% (v/v) rat normal serum. Bound nEgAgB was detected with a 1/100 dilution of a rabbit antiserum anti-EgAgB8/1 generously donated by Betina Córsico and Gisella Franchini (INIBIOLP, Facultad de Ciencias Médicas, Universidad Nacional de La Plata, Argentina). In parallel, a normal rabbit antiserum was used for controlling non-specific binding. After washing, BMDC-bound immune complexes were detected by incubation with anti-rabbit IgG-Alexa 488 and anti-CD11c antibodies. For each cell type (WT, *Tlr4*^-/-^ and *Tlr2*^-/-^) a binding index was calculated to quantify nEgAgB binding. This index was determined as the increase of the Alexa488 fluorescence intensity (FI, geometric mean) on CD11c+ cells relative to the Veh (control) and expressed as a percentage.

### Analysis of EgAgB effects on cell surface CD14, TLR4 and TLR4 dimerization/endocytosis

BMDC (0.4 x 10^6^ cells/well, in duplicates) were incubated with nEgAgB or rEgAgB (1 or 10 µg/mL) or Veh in the presence or absence of LPS (10 ng/mL), at 37°C in a humidified atmosphere with 5% CO_2_ (v/v). After 2 hs, cells were collected by gentle pipetting and transferred to a V-bottom plate. Staining of cell surface receptors was performed following conventional protocols with anti-CD11c-PECy7, anti-CD14-FITC, anti-TLR4/MD2-PE and anti-TLR4-APC (all from BioLegend). Cells were acquired on a FACSCanto II cytometer, and analysed using the FlowJoTM package (Version 7.6.2), gating on CD11c^+^ events. For data analysis, the relative expression levels of CD14 and total TLR4 (measured with the anti-TLR4-APC that recognizes both monomeric and dimeric TLR4) were estimated as the median of the fluorescence intensity values (MFI, geometric mean) normalized to the unstimulated cell condition. The percentage of TLR4 dimerization/endocytosis was determined using the anti-TLR4/MD2-PE (recognises only the monomeric TLR4/MD2 complex) and calculated as 100% -[(FI of sample (geometric mean)/FI of unstimulated cell condition (geometric mean)) x 100], as described previously (27).

### Interference with LPS binding to cells

EgAgB’s ability to interfere with LPS for binding to BMDC or THP-1 macrophages was analysed by flow cytometry. All incubations and washing steps were performed in the FACS buffer. Cells (0.2 x 10^6^) were seeded in conical-bottom plates and incubated with 50 μL of EgAgB, hHDL (both 1-200 μg/mL), ovalbumin (OVA, 200 μg/mL as a control protein) or vehicle (Veh) for 30 min at 37°C. Subsequently, 50 μL of LPS conjugated to the fluorochrome Alexa Fluor 488 (5 μg/mL, from *E. coli* O55:B5, Molecular Probes) was added and incubated 30 min at 37°C. After incubation, cells were washed twice, acquired on a FACSCalibur cytometer (BD Biosciences) and analysed using the FlowJoTM package (Version 7.6.2). The fluorescence intensity (geometric mean) of all samples was corrected by the Veh. LPS binding was expressed as the average of the fluorescence intensity of samples relative to the average of the intensity corresponding to the LPS-Alexa-488 (100%).

### Analysis of EgAgB-LPS interaction by DLS

EgAgB or hHDL (both at 1 mg/mL) were mixed with an equal volume (35 µL) of LPS (1 mg/mL) and subsequently analysed by DLS as described above. For comparison, EgAgB, hHDL, and LPS were mixed with the vehicle control (PBS_EB_) and analysed under identical conditions. The mean hydrodynamic diameter of the particle size distribution (Z-average) was calculated.

### ELISA assay for LPS binding activity

EgAgB ability to bind LPS was analysed based on the protocol previously reported (22). To that end, nEgAgB, hHDL/hHDL_3_ as positive controls or OVA as an irrelevant control protein (10 μg/mL, 100 μL/well) were immobilized in high-binding ELISA 96-well plates (Maxisorp, Nunc) overnight at 4°C. Specifically, the hHDL_3_ fraction (density range from 1,125 g/mL-1,210 g/mL) was freshly obtained from pooled plasma by sequential flotation ultracentrifugation as described previously (25). In parallel, wells incubated with PBS were prepared to evaluate nonspecific interactions with the blocker (blocker control). Blocking was done with PBS-BSA 1% for 1.5 h at room temperature (RT). Various concentrations of LPS conjugated to biotin at its inner core (0, 10, 100, and 1000 ng/mL, from *E. coli* O111:B4, InvivoGen #tlrl-lpsbiot) were preincubated with or without hLBP (human LBP, 0.1 μg/mL, R&D Systems #6445-LP) for 1 h at RT and then incubated in the plates for 45 min at 37°C. Subsequently, streptavidin-HRP (Sigma #5512) was added for another 45 min at 37°C. Between incubations, five washes with PBS Tween 0.05% were performed, except after sensitization when two washes were done. For development, acetate 0.1 M containing TMB and H_2_O_2_ was used and the reaction was stopped with H_2_SO_4_ 1 M. The absorbance was quantified at 450 nm, correcting with the absorbance at 560 nm. The values obtained for samples were corrected by subtraction of their corresponding controls of non-specific interactions, and then graphed against the concentration of LPS-biotin using GraphPad software (10.4.1 Version). rEgAgB could not be assessed since it contains a strep-tag motif necessary for its purification that interferes with the colour developing step.

### *In silico* analysis of EgAgB8/1 interaction with LPS

The SMILES structures corresponding to the R2 and R3 core domains of lipopolysaccharide (LPS) were retrieved from the BioCyc database, under compound identifiers CPD-21363 and CPD-21361, respectively. The SMILES structure of lipid A was obtained from the ChEBI database (ChEBI ID: CHEBI:134256). The three-dimensional (PDB) structures of R2, R3, and lipid A were generated from their respective SMILES strings using the SwissParam tool (28). The structure of the O-antigen domain was also derived from ChEBI (ChEBI ID: CHEBI:89981), corresponding to a full LPS molecule. The SMILES file was downloaded, and the O-antigen region was manually isolated by truncating the non-relevant portions of the molecule. The resulting fragment was converted to PDB format using Open Babel (29). All generated PDB files were geometry-optimized using IQmol (version 2.15), employing the MMFF94 force field. The primary sequence of the *Echinococcus granulosus* apolipoprotein EgAgB8/1 was obtained from the UniProt database (UniProt ID: U6JQF4). The corresponding three-dimensional structure was obtained from the theoretical model predicted by AlphaFold (30). Molecular docking simulations were performed using the HADDOCK web server (version 2.4) (31), following the standard protein–ligand docking protocol. For each EgAgB1-LPS domain pair, the top-scoring model from the most populated cluster was selected for further analysis. Protein-ligand interactions were characterized using the Protein-Ligand Interaction Profiler (PLIP) (32), and three-dimensional visualization and inspection of docking poses were conducted in PyMOL (Schrödinger, LLC; version 2.5).

### Data analysis

The number of independent experiments and analytical repetitions is provided in each figure legend. For graphical presentation, some data were normalized to either the control group or the LPS response, as specified in figure legends. Statistical analyses were performed on the raw data: for assays using the THP-1 cell line, we assume normality and homoscedasticity, and a two-way ANOVA with Tukey’s post hoc test was applied (using GraphPad Prism 9.4.1); for BMDC experiments, as we observed more dispersion between experiments and normality and homoscedasticity could not be checked, the two-way non-parametric Friedman test with Bonferroni correction for multiple comparisons was applied. The statistical significance is indicated in graphs by symbols (i.e. *, $, #) and explained in each figure legend.

## Results

### nEgAgB and rEgAgB preparations exhibited opposite effects on BMDC: they induced a modest activation on their own but interfered with LPS-driven activation

To obtain highly pure preparations, both nEgAgB and rEgAgB were purified using a final immunoaffinity chromatography step employing an anti-EgAgB8/1 nanobody developed by our research group (14). As observed in the referenced study, DLS analysis of EgAgB preparations showed multimodal size distributions (weighted by intensity) corresponding to principal components (weighted by volume) with an average D_H_ of 13.4 ± 1.6 nm for nEgAgB preparations and 17.5 ± 2.0 nm for rEgAgB (Supplementary Fig. 1 and 3 for nEgAgB and rEgAgB forms, respectively). The molecular size and protein profile obtained by SDS-PAGE and Western Blot analysis of the resulting preparations are illustrated in Supplementary Fig. 3, being consistent with those previously described (14). Based on the LAL assay, LPS content in most nEgAgB and rEgAgB preparations was comparable to those previously reported (≤ 0.02 ng/µg protein, (14)). Interestingly, while our observations in macrophages indicated that concentrations of up to 10 µg/mL of nEgAgB/rEgAgB did not induce cell activation (14), BMDC exposure to EgAgB preparations as the sole stimulus led to modest yet significant secretion of the inflammatory cytokines IL-6 and IL-12p40 (Fig. 1A,B and 1C,D, respectively), albeit with IL-12p40 being secreted to a lesser extent, as well as ·NO production (Fig. 1G,H). These responses were not accompanied by the secretion of IFN-β (Fig. 1E,F). Remarkably, the activating effect of EgAgB on BMDC did not hinder its ability to modulate LPS-driven cellular responses in a dose-dependent manner. Both native and recombinant EgAgB preparations limited IL-6, IL-12p40 and IFN-β secretion and the generation of ·NO in LPS-activated BMDC (Fig. 1). Consistent with our previous findings in macrophages (11), nEgAgB and rEgAgB only caused modest effects on costimulatory molecules despite their impact on IL-6 and IL-12p40 secretion; they slightly reduced CD40 expression while did not affect CD86 expression (Fig. 2A,C and 2B,D, respectively).

**Figure 1.**
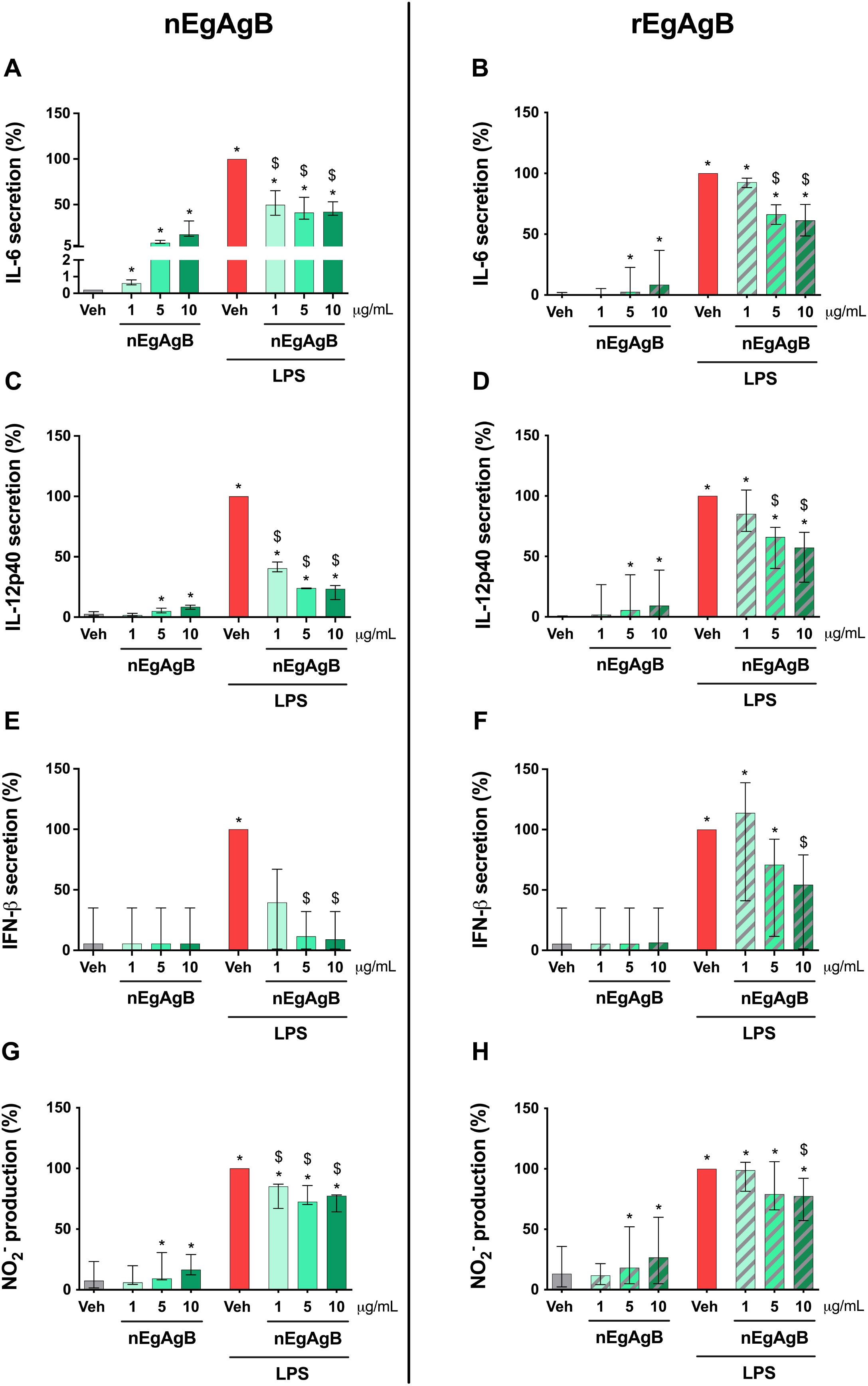
Effects of nEgAgB and rEgAgB on the cytokine secretion and nitrite formation by BMDC in the absence or presence of LPS. BMDC (0.4×10^6^) were stimulated with nEgAgB/rEgAgB (1, 5 or 10 µg/mL) or PBS_EBAb_ (vehicle, Veh) in absence or presence of LPS (10 ng/mL). After 18 hs, the culture supernatant was collected to measure cytokines secretion by ELISA or nitrite by Griess. The levels of IL-6 (A, B), IL-12p40 (C, D), IFN-β (E, F) and nitric oxide (·NO) generation (G, H) are plotted as the median and range of data normalised to the LPS condition (set as 100%). Data correspond to three or four independent experiments with analytical duplicates. Significant differences with Veh or Veh+LPS are indicated with * and $, respectively (Friedman test with the Bonferroni correction for multiple comparisons, p < 0.05).

**Figure 2.**
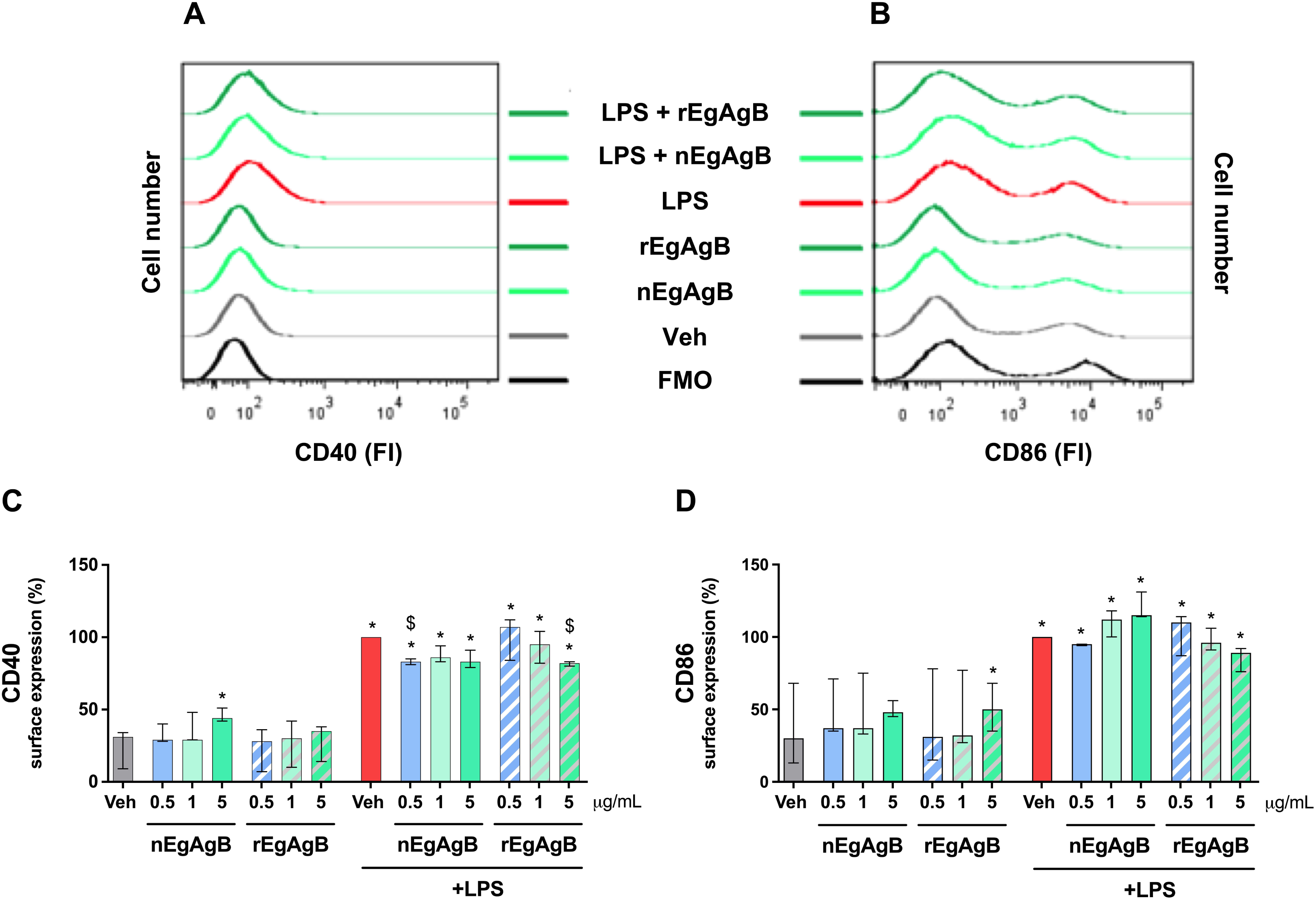
Effects of nEgAgB and rEgAgB on the surface expression of costimulatory molecules by BMDC in the absence or presence of LPS. BMDC (0.4×106) were stimulated with nEgAgB/rEgAgB (0,5 1, 5 µg/mL) or Veh in absence or presence of LPS (10 ng/mL). After 18 hs, the cells were recovered to measure the surface expression of CD40 and CD86 costimulatory molecules (within live CD11c+ cells) by flow cytometry. Representative histograms of the fluorescence intensity (FI) obtained for CD40 (A) and CD86 (B) surface expression analysis are shown. Bar graphs correspond to CD40 (C) and CD86 (D) expression, presented as the median and range of FI values normalised to the Veh+LPS condition (set as 100%). Data correspond to three independent experiments with analytical duplicates. Significant differences with Veh or Veh+LPS are indicated with * and $, respectively (Friedman test with Bonferroni correction for multiple comparisons, p < 0.05).

### BMDC activation by EgAgB preparations was partially inhibited by polymyxin B and dependent on TLR4 signalling

Despite efforts to avoid LPS contamination during purification, some EgAgB preparations contained trace levels of LPS according to the LAL assay, suggesting that the observed EgAgB activating effect on BMDC was due to LPS. To examine this, we firstly selected the nEgAgB/rEgAgB batches showing the highest endotoxin level (0.02–0.09 ng/µg protein) and used polymyxin B for LPS neutralization (33). At 10 µg/mL, polymyxin B effectively inhibited IL-6 and IL-12p40 secretion by BMDC stimulated with 10 ng/mL LPS, while it did not affect the response to 60 ng/mL Pam3CSK4, serving as positive and negative controls, respectively; these controls validate the polymyxin B efficacy under the assessed conditions. Given the estimated LPS content in EgAgB preparations (<0.09 ng/mL for tested concentrations), we anticipated that 10 µg/mL polymyxin B would inhibit any putative LPS activity in these preparations. However, polymyxin B partially reduced IL-6 and IL-12p40 secretion driven by nEgAgB and rEgAgB (Fig. 3A,B).

**Figure 3.**
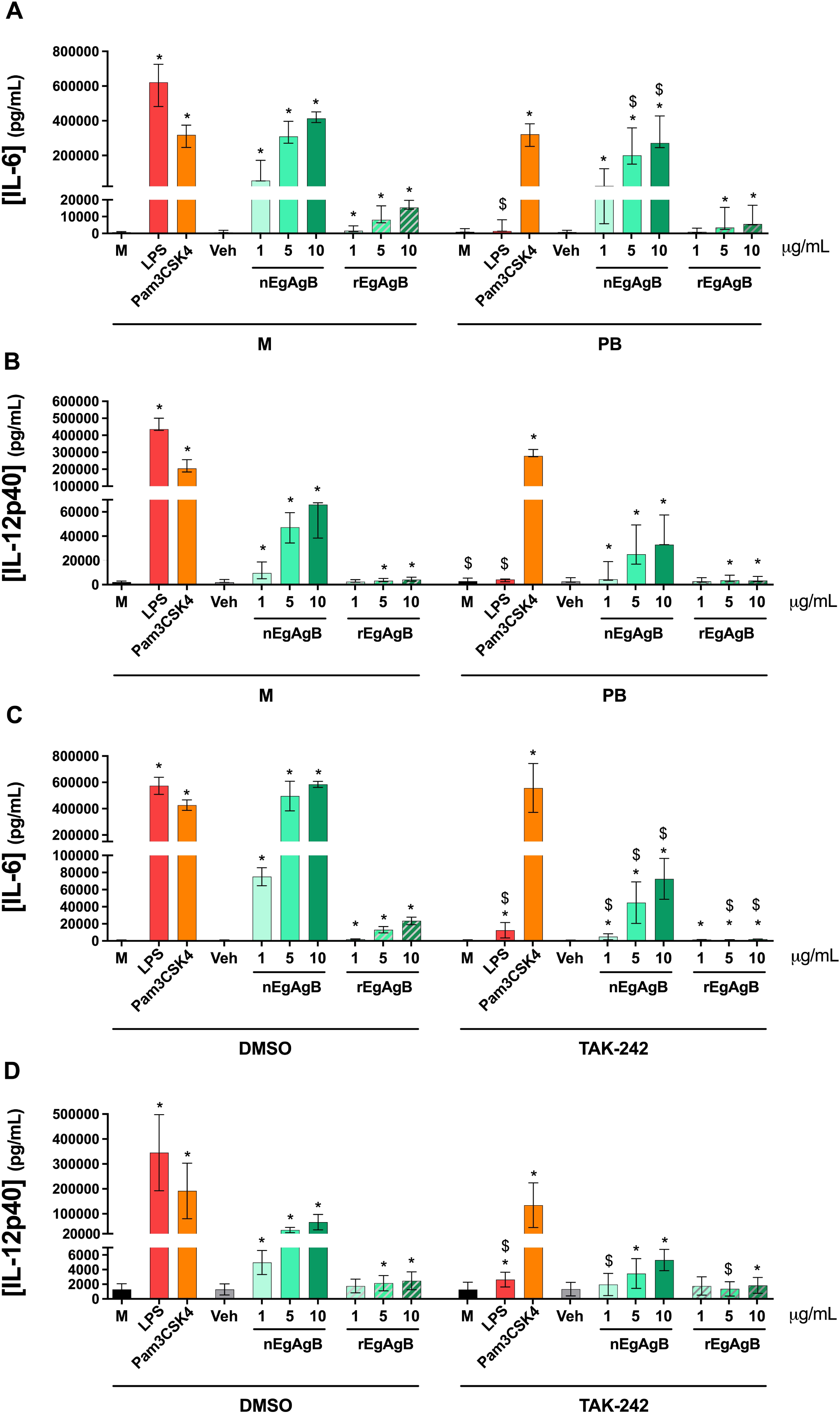
BMDC activation by nEgAgB and rEgAgB was nearly abolished by the TLR4 inhibitor TAK-242, but not by polymyxin. **B.** In panels A and B, BMDC (0.4×10^6^) were treated with polymyxin B (PB, 10 µg/mL) or cultured medium (control) for 2 hs and subsequently stimulated with nEgAgB/rEgAgB (1, 5 or 10 µg/mL), LPS (10 ng/mL), Pam3CSK4 (60 ng/mL). Culture medium (M) and PBS_EBAb_ (Veh) served as stimulation controls. After 18 hs, cytokine secretion was measured in the culture supernatants by ELISA. The levels of IL-6 (A) and IL-12p40 (B) are plotted as the median and range of data from three independent experiments with analytical duplicates. Significant differences with the corresponding stimulation control (M or Veh) are indicated with *, and significant differences between selected conditions are indicated with $ (Friedman test with Bonferroni correction for multiple comparisons, p < 0.05). In panels C and D, BMDC (0.4×10^6^) were pretreated with TAK-242 (10 µM) or DMSO (control pre-treatment) for 1 h and subsequently stimulated with nEgAgB/rEgAgB (1, 5 or 10 µg/mL) or PBS_EBAb_ (Veh stimulation control). After 18 hs, cytokine secretion was measured in the culture supernatants by ELISA. The levels of IL-6 (A), IL-12p40 (B) are presented as the median and range of data from two independent experiments with analytical duplicates. Significant differences compared to the corresponding stimulation control (M or Veh) within pretreated or control BMDC are indicated with *, and significant differences between equivalent stimulation conditions of BMDC pretreated with TAK-242 or DMSO are indicated with $ (Friedman test with Bonferroni correction for multiple comparisons, p < 0.05). Note that for both studies we used the nEgAgB with higher endotoxin activity according to LAL (0.02–0.09 ng/µg protein).

To explore whether the cytokine response elicited by EgAgB preparations in BMDC was dependent on TLR4 signalling, we employed the pharmacological inhibitor TAK-242, which specifically binds to a cytoplasmic domain of TLR4 and interferes with the receptor signalling. Pretreatment of BMDC with TAK-242 inhibited IL-6 (97.8%) and IL-12p40 (99.2%) secretion induced by LPS (10 ng/mL), but not by Pam3CSK4 (60 ng/mL) (Fig. 3C,D). Interestingly, the dose-dependent cytokine response induced by nEgAgB and rEgAgB was inhibited following TAK-242 pretreatment (Fig. 3C,D); however, a residual cytokine secretion (mainly IL-6) was detected.

To confirm the involvement of TLR4 in BMDC activation by EgAgB, we used BMDC from WT, *Tlr2*^-/-,^ and *Tlr4*^-/-^ mice and nEgAgB/rEgAgB batches with lower LPS content (≤ 0.02 ng/µg protein). Consistent with our previous results, both nEgAgB and rEgAgB induced dose-dependent IL-6 secretion by WT BMDC (Fig. 4A,B), but IL-12p40 secretion was undetectable (a trend towards an increase was observed only for nEgAgB, which may be due to the lower LPS content, Fig. 4C). The IL-6 responses induced by nEgAgB and rEgAgB were comparable between WT and *Tlr2^-/-^*BMDC, but almost null in *Tlr4*^-/-^ BMDC, which were unresponsive to rEgAgB and secreted low levels of IL-6 at higher nEgAgB concentrations (≥ 5 µg/mL). These findings suggested that the ability of nEgAgB and rEgAgB lipoproteins to activate BMDC was primarily via TLR4, despite a minor contribution of TLR4-independent mechanisms may play a role in the case of nEgAgB.

**Figure 4.**
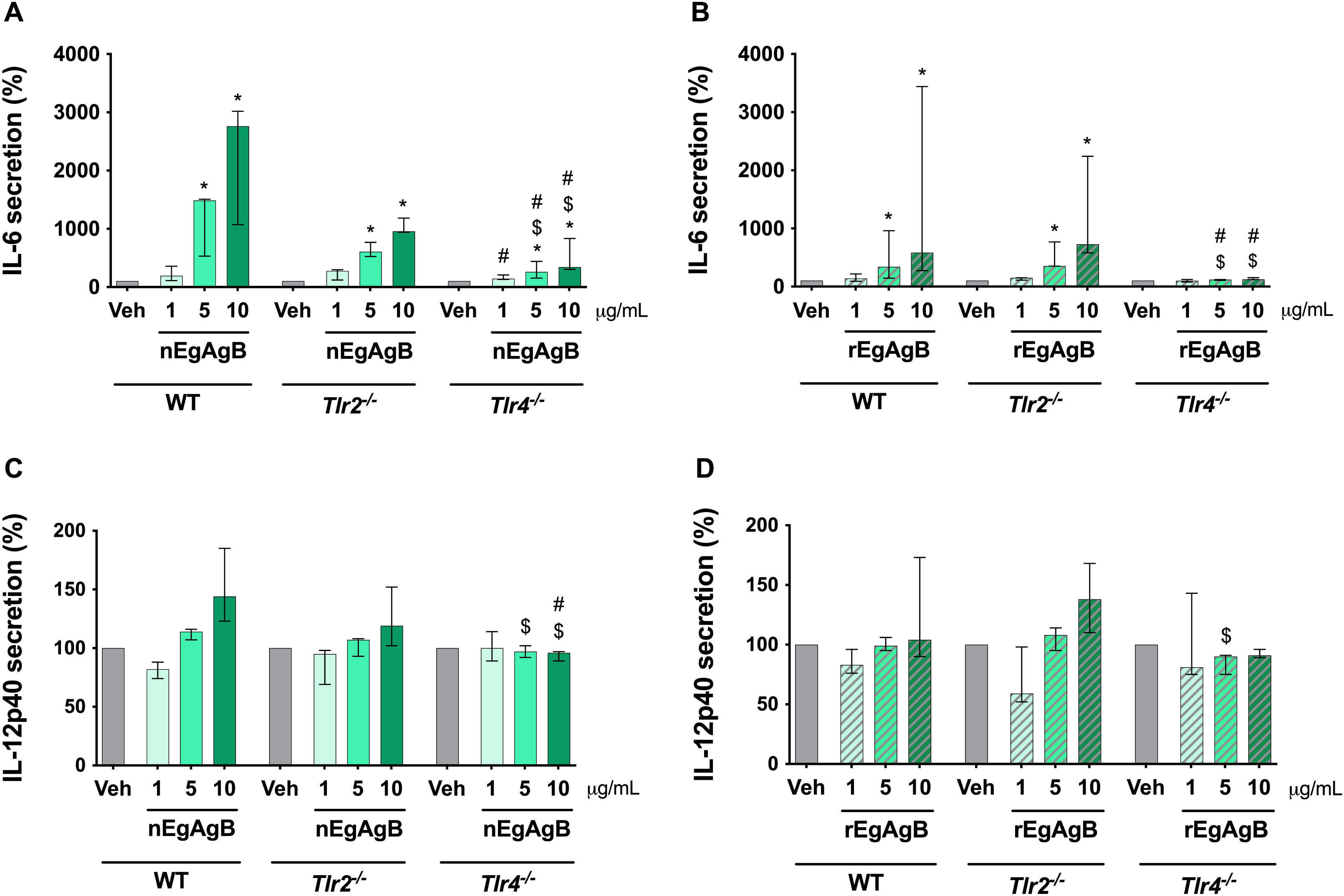
The cytokine response induced by nEgAgB and rEgAgB in BMDC depended on TLR4. BMDC (0.4×10^6^) derived from WT, *Tlr4*^-/-^ and *Tlr2*^-/-^mice were stimulated with nEgAgB/rEgAgB (1, 5 or 10 µg/mL) or PBS_EBAb_ (Veh). After 18 hs, cytokine secretion was measured in the culture supernatants by ELISA. The levels of IL-6 (A, B) and IL-12p40 (C, D) are presented as the median and range of data normalised to the response of WT BMDC stimulated with 10 µg/mL of EgAgB (set as 100%). Data correspond to four independent experiments with analytical duplicates. Significant differences are indicated according to: comparisons with Veh within the same strain are marked with *, comparisons between equivalent stimulation conditions in *Tlr2*^-/-^ and *Tlr4*^-/-^⁻ BMDC with WT BMDC are marked with $, and comparisons between equivalent stimulation in *Tlr2*^-/-^ and *Tlr4*^-/-^ BMDC are marked with # (Friedman test with Bonferroni correction for multiple comparisons, p < 0.05).

### EgAgB bound to BMDC in a TLR4-independent manner

In addition to LPS, TLR4 recognizes various endogenous ligands, including minimally modified lipoproteins (34, 35) and saturated fatty acids (36). Besides, EgAgB was found to bind to various myeloid cell types (12, 37). To determine whether EgAgB utilizes TLR4 for cell binding and signalling, we measured EgAgB binding to BMDC from WT, *Tlr2*^-/-^, and *Tlr4*^-/-^ mice. The results showed that nEgAgB binds to BMDC independently of the presence of TLR4 and TLR2 (Fig. 5). Collectively, the data suggested that TLR4-dependent activation of BMDC by EgAgB preparations likely results from LPS carried by EgAgB lipoproteins rather than from a direct EgAgB-TLR4 interaction.

**Figure 5.**
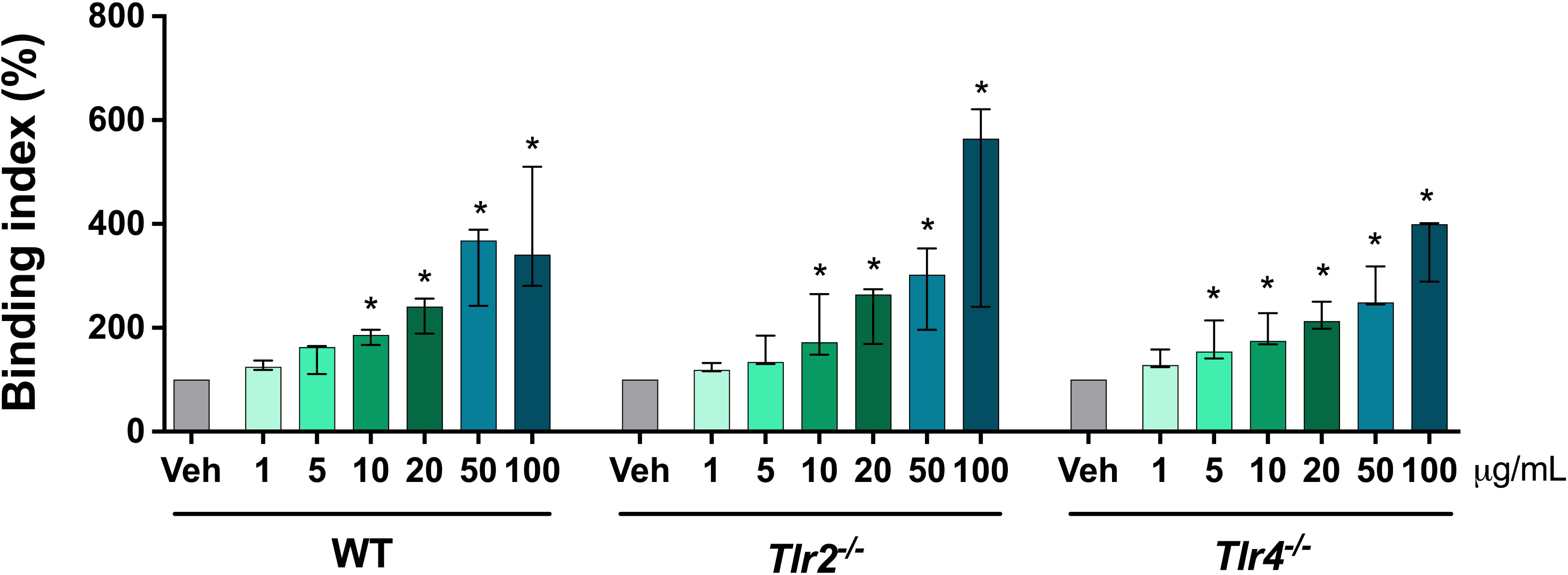
nEgAgB and rEgAgB bound BMDC in a TLR4-independent manner. BMDC (0.2×10^6^) derived from WT, *Tlr4*^-/-^ and *Tlr2*^-/-^ mice were incubated with nEgAgB (1-100 µg/mL) or PBS_EBAb_ (Veh) in the binding buffer at 4°C. After 1 h, cells were washed, Fc receptors were blocked, and bound nEgAgB was detected using an anti-EgAgB8/1 or a normal rabbit antiserum (non-specific binding control) followed by an anti-rabbit-IgG antibody conjugated to Alexa 488. nEgAgB binding to BMDC (gated as CD11c+ cells) is plotted as the binding index of data from three independent experiments with analytical duplicates. Significant differences compared to Veh within each BMDC type are indicated with * (Friedman test with Bonferroni correction for multiple comparisons, p < 0.05). No significant differences were observed in the binding index of nEgAgB at equal concentrations across BMDC derived from the three mice strains.

### EgAgB interfered with LPS-triggered TLR4 dimerization/endocytosis

Despite the activating capacity of EgAgB preparations, both nEgAgB and rEgAgB were found to suppress LPS-driven activation of BMDC. To investigate the molecular mechanisms underlying this modulation, we examined whether EgAgB could alter LPS-mediated changes in cell surface CD14, a co-receptor that facilitates LPS binding to MD2, which is a crucial accessory molecule for TLR4 activation (38). As shown in Fig. 6A, LPS treatment induced a modest increase in cell surface CD14 expression, which was counteracted by the presence of nEgAgB (10 µg/mL), but not by rEgAgB. Additionally, LPS stimulation caused a trend towards a decrease in cell surface total TLR4, but this trend was not observed in the presence of nEgAgB or rEgAgB (Fig. 6B)). A decreased cell surface TLR4 expression is expected as a consequence of LPS-driven TLR4 dimerization and subsequent endocytosis. To further explore this, we assessed TLR4 dimerization/endocytosis using a mAb targeting the TLR4/MD2 complex (MTS510), which indirectly allows its measurement by detecting the loss of monomeric TLR4/MD2 from the cell surface (27). BMDC exposure to LPS led to a significant TLR4 dimerization/endocytosis after 120 minutes (Fig. 6C). Notably, in the presence of both nEgAgB and rEgAgB this LPS-driven process was reduced, although the rEgAgB’s effect did not reach statistical significance. Our findings suggest that the modulatory effects of EgAgB on LPS-induced BMDC activation are linked to molecular interactions established by these parasite lipoproteins, which disrupt the ability of LPS to fully engage and activate the initial steps of the TLR4 signalling pathway.

**Figure 6.**
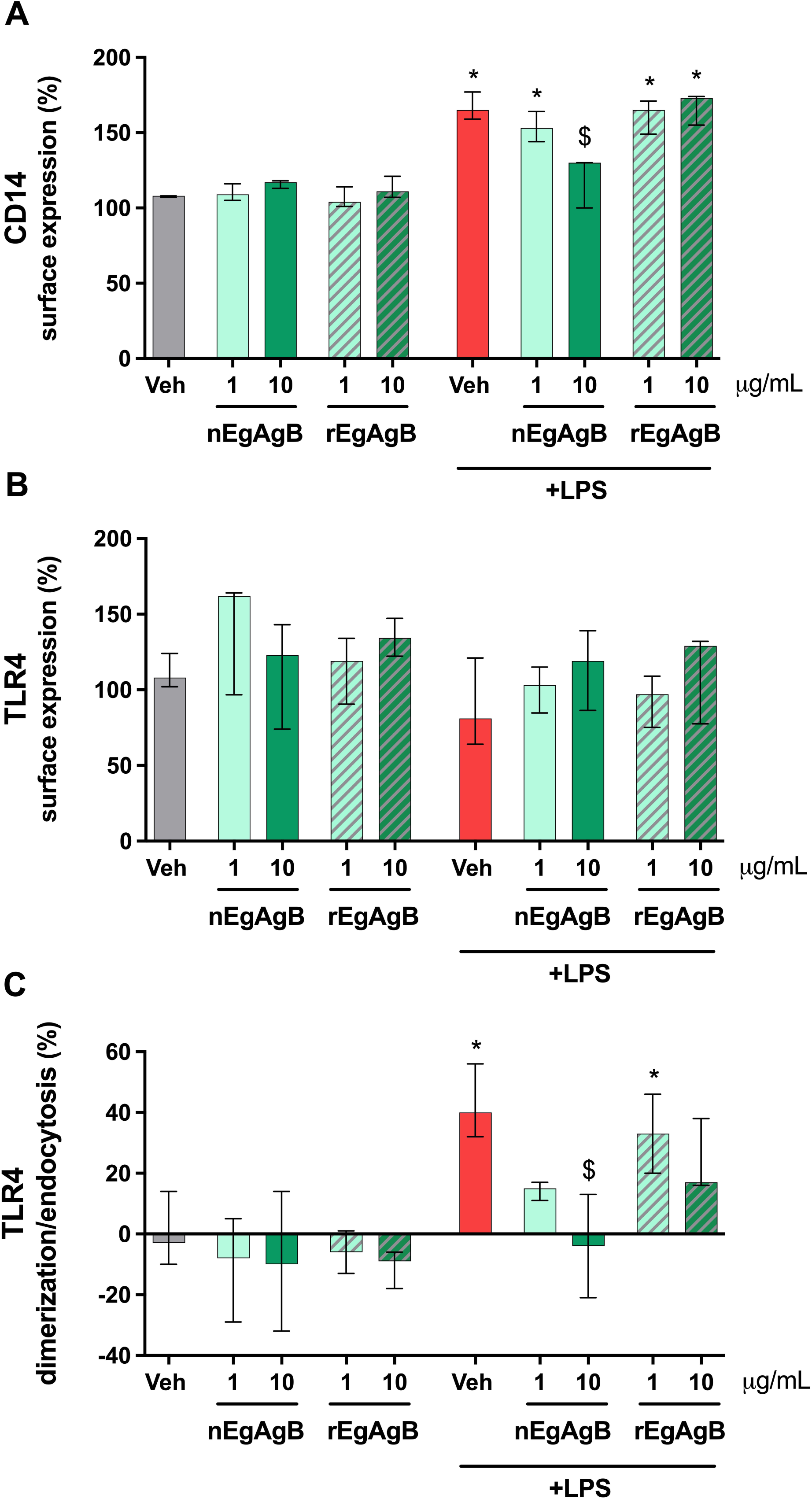
nEgAgB and rEgAgB inhibited LPS-triggered TLR4 dimerization/endocytosis in BMDC. BMDC (0.4×10^6^) were stimulated with nEgAgB/rEgAgB (1 or 10 µg/mL) or PBS_EBAb_ (Veh) in the absence or presence of LPS (10 ng/mL). After culture for 2 hs, cells were recovered and the surface expression of CD14, total TLR4 and monomeric TLR4/MD2 was measured by flow cytometry. The surface expression of CD14 (A), TLR4 (B) and levels of TLR4 dimerization/endocytosis (C) on BMDC (gated as CD11c+ cells) are plotted as the median and range of data normalised to the unstimulated cell condition (set as 100%). Data correspond to three independent experiments with analytical duplicates. Significant differences from comparisons with Veh or Veh+LPS are indicated with * and $, respectively (Friedman test with Bonferroni correction for multiple comparisons, p < 0.05).

### EgAgB interfered with LPS binding to myeloid cells, similar to the action of hHDL

The inhibition of LPS-triggered TLR4 dimerization/endocytosis by EgAgB may involve interference with LPS binding to TLR4^+^ cells. To test this hypothesis, we conducted an inhibition binding assay employing LPS-Alexa-488 and PMA-differentiated THP-1 macrophages or BMDC as models of TLR4^+^ myeloid cells. Additionally, two control groups were included; a positive one with hHDL, an LPS scavenger macromolecule (22), and a negative control with OVA as an irrelevant protein. Pre-incubation with OVA showed no effects on LPS binding to THP-1 macrophages, while the presence of 200 μg/mL hHDL reduced the binding index by approximately 23% (Fig. 7). For testing EgAgB in this assay, we used nEgAgB preparations purified from HF by immunoaffinity, with or without prior ultracentrifugation. Both nEgAgB preparations showed a dose-dependent ability to interfere with LPS binding to THP-1 macrophages, exhibiting higher inhibitory activity than hHDL (around 64% at 200 μg/mL, Fig. 7). nEgAgB also showed interference activity with LPS binding to BMDC (Supplementary Fig. 4). Taking into account that EgAgB binding to BMDC was TLR4-independent, EgAgB’s immunomodulatory effects on LPS-driven myeloid cell activation likely involve its interaction with LPS.

**Figure 7.**
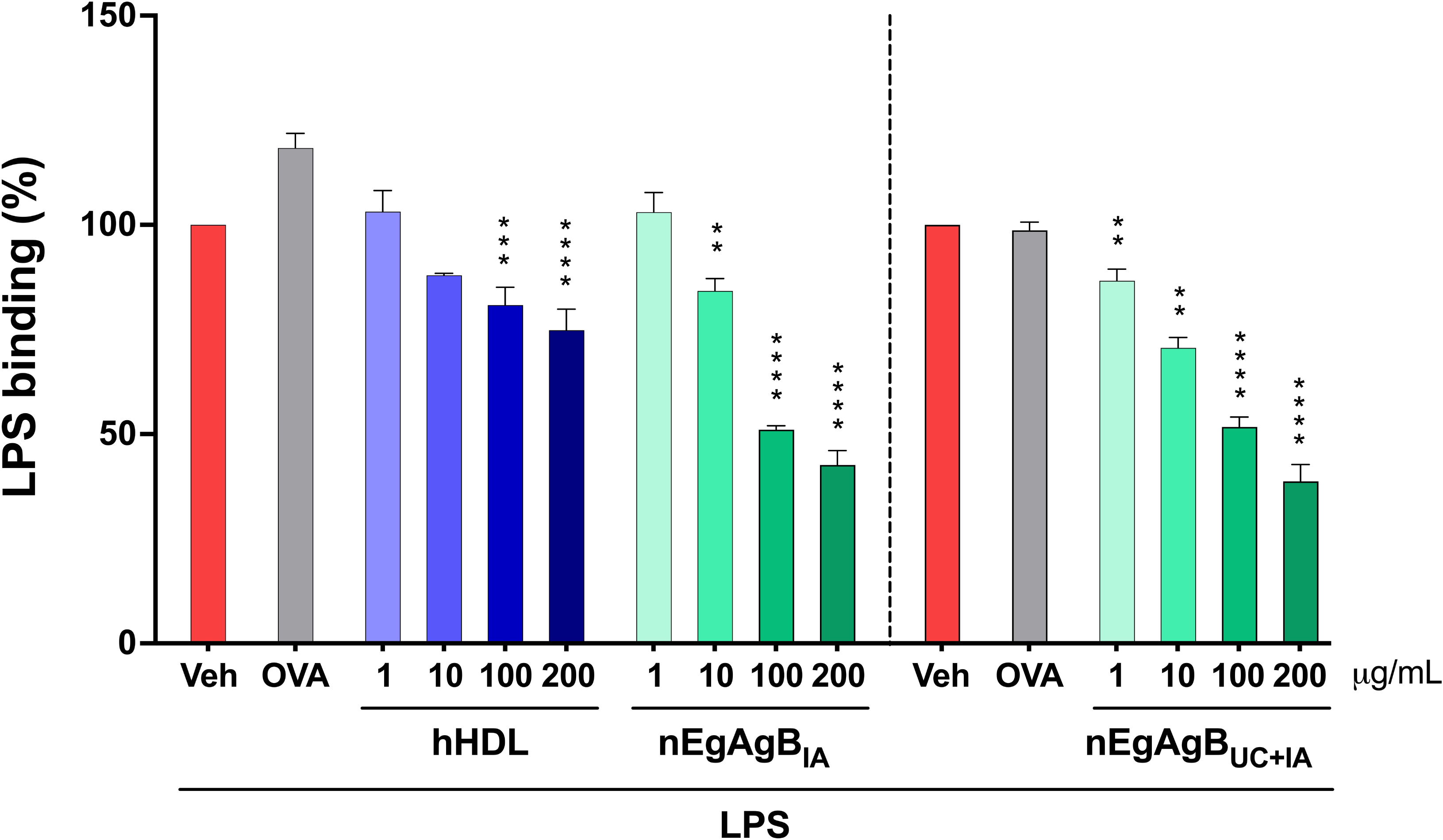
nEgAgB competed with LPS for binding to THP-1 macrophages, as hHDL does. THP-1 macrophages (0.2×10^6^) were incubated for 30 min with nEgAgB, hHDL (both 1-200 μg/mL), OVA (200 μg/mL) or PBS_EBAb_ (Veh control), followed by incubation with LPS-Alexa-488. Two preparations of nEgAgB were assessed: one purified including the ultracentrifugation step (nEgAgB_UC+IA_) and one without this step (nEgAgB_IA_). After washing, cell fluorescence was measured by flow cytometry. LPS binding to THP-1 macrophages is presented as the mean ± SD of data normalised to the LPS condition in the absence of competitive molecules (Veh control). Data correspond to three independent experiments with analytical duplicates. Significant differences compared to Veh or between the selected conditions are indicated with * and $, respectively (two-way ANOVA and Tukey’s test, p < 0.05). Note that no differences were observed among nEgAgB_UC+IA_ and nEgAgB_IA_ ability to compete with LPS binding to THP-1 macrophages.

### EgAgB showed a superior ability to modulate LPS-induced cytokine secretion by BMDC than hHDL

Because hHDL has been described as a scavenger molecule of some PAMPs like LPS and LTA (21), we compared the ability of EgAgB and hHDL to modulate BMDC activation induced by these TLR agonists. In the assayed conditions, concentrations of 100 μg/mL of hHDL were needed to cause a modest inhibition of LPS-induced IL-6 secretion in BMDC (Fig. 8A). In parallel, nEgAgB and rEgAgB showed higher inhibitory activity compared to hHDL, achieving approximately 50% reduction in the IL-6 response at just 10 μg/mL, an inhibition level unattainable even with 100 μg/mL of hHDL (Fig. 8B). In THP-1 macrophages, hHDL caused a robust inhibition of LPS-driven IL-6 secretion, but again 10-fold higher concentrations of hHDL than nEgAgB were needed to achieve a similar percentage of inhibition (Supplementary Fig. 5). Additionally, nEgAgB and rEgAgB, but not hHDL, significantly suppressed IL-12p40 secretion in LPS-activated BMDC (Fig. 8C, D). In contrast, when LTA was used as a TLR2 agonist, hHDL, but not nEgAgB and rEgAgB, diminished the secretion of IL-6 (Fig. 8E,F) and IL-12p40 (Fig. 8G,H). Paradoxically, both EgAgB preparations increased LTA-induced IL-6 and IL-12p40 responses. Overall, these findings suggest a selective ability of EgAgB to interfere with LPS-mediated activation.

**Figure 8.**
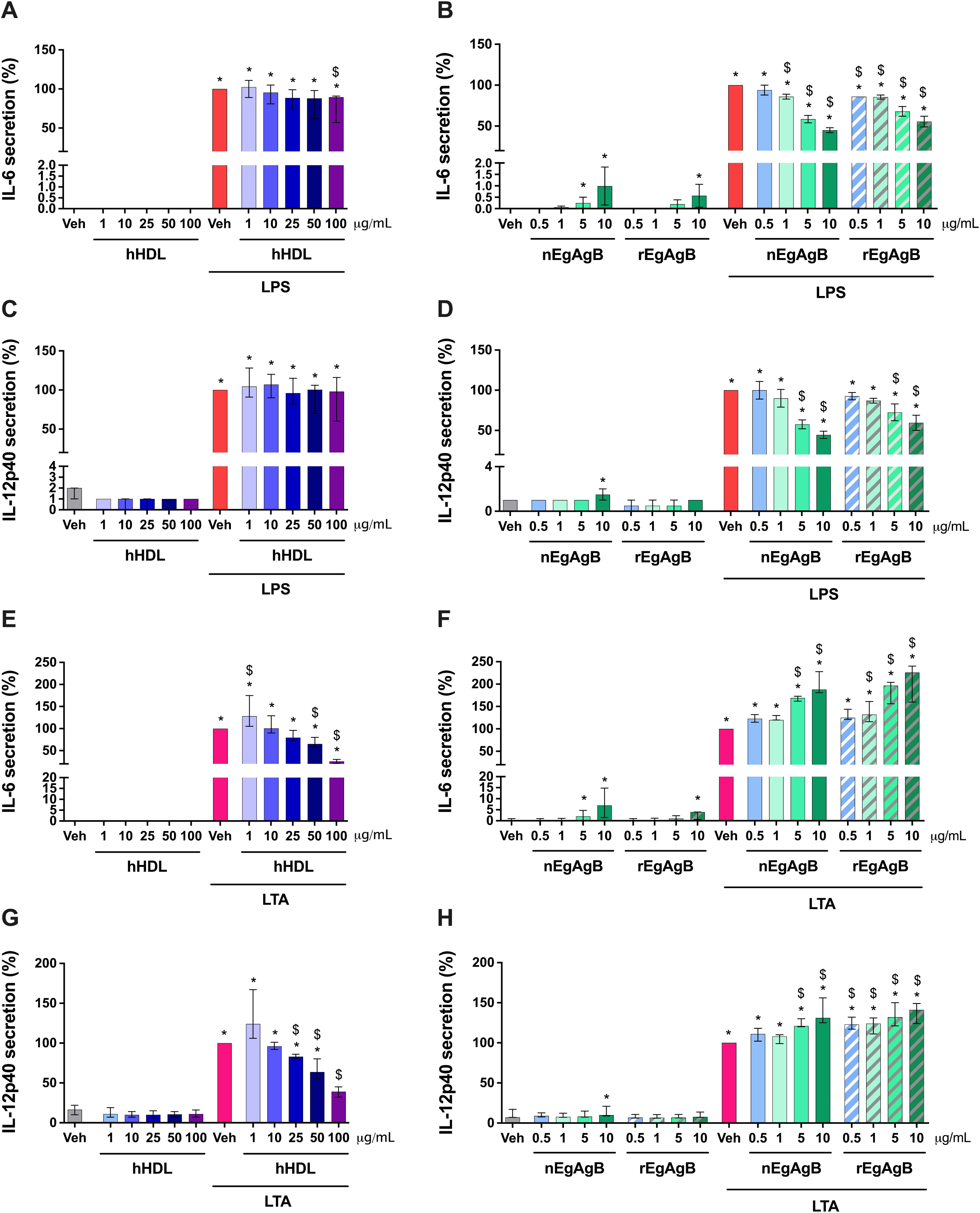
nEgAgB/rEgAgB, unlike hHDL, inhibited LPS-induced but not LTA-induced activation of BMDC. BMDC (0.4×10^6^) were stimulated with nEgAgB (0.5, 1, 5 or 10 µg/mL), hHDL (1-100 µg/mL) and PBS_EBAb_ (Veh control) in the absence or presence of LPS (10 ng/mL) or LTA (0.5 µg/mL). After 18 hs, cytokine secretion was measured in the culture supernatants by ELISA. Graphs show the levels of IL-6 and IL-12p40 as the median and range of data normalised to the corresponding LPS+Veh or LTA+Veh condition (both set as 100%). Data correspond to three independent experiments with analytical duplicates. Significant differences compared to Veh or LPS/LTA+Veh are indicated with * and $, respectively (Friedman test with Bonferroni correction for multiple comparisons, p < 0.05).

### EgAgB demonstrates LPS-binding capacity like hHDL and hHDL_3_

Our findings suggested that EgAgB’s modulation of LPS-triggered activation of BMDC could stem from EgAgB binding to LPS, thereby interfering with its interaction with the TLR4/MD2 complex. To study the potential of EgAgB to bind LPS, we initially analysed changes in the size distribution of EgAgB lipoprotein particles upon LPS addition using DLS, with hHDL included for comparison. Results showed that LPS modified the size distribution of EgAgB in solution by promoting both a partial shift of the predominant population to smaller species and an increase in the abundance of supramolecular assemblies which remain scarce (Supplementary Fig. 6). This alteration in the multimodal size distribution (weighted by intensity) of EgAgB was not as noticeable for hHDL (Supplementary Fig. 6). Unfortunately, despite using similar mass concentrations of both LPS and EgAgB, we cannot estimate the molar concentration of each species and figure out the relevance of the observed shifts in size. Since LPS shows a size profile superimposed with one of the populations observed in the complex profile of EgAgB (Supplementary Fig. 6), we used the intensity weighted Z-average diameter -as an independent magnitude describing the overall mixture profile-to evaluate the effect of LPS addition and compare its significance with hHDL. Interestingly, the Z-average of the particles present in both native and recombinant EgAgB-LPS mixtures was higher than that corresponding to EgAgB alone (Fig. 9A), suggesting that addition of LPS led to the emergence or stabilization of larger assemblies as a consequence of an EgAgB-LPS interaction. In contrast, no increase in Z-average was detected by adding LPS to hHDL (Fig. 9A).

**Figure 9.**
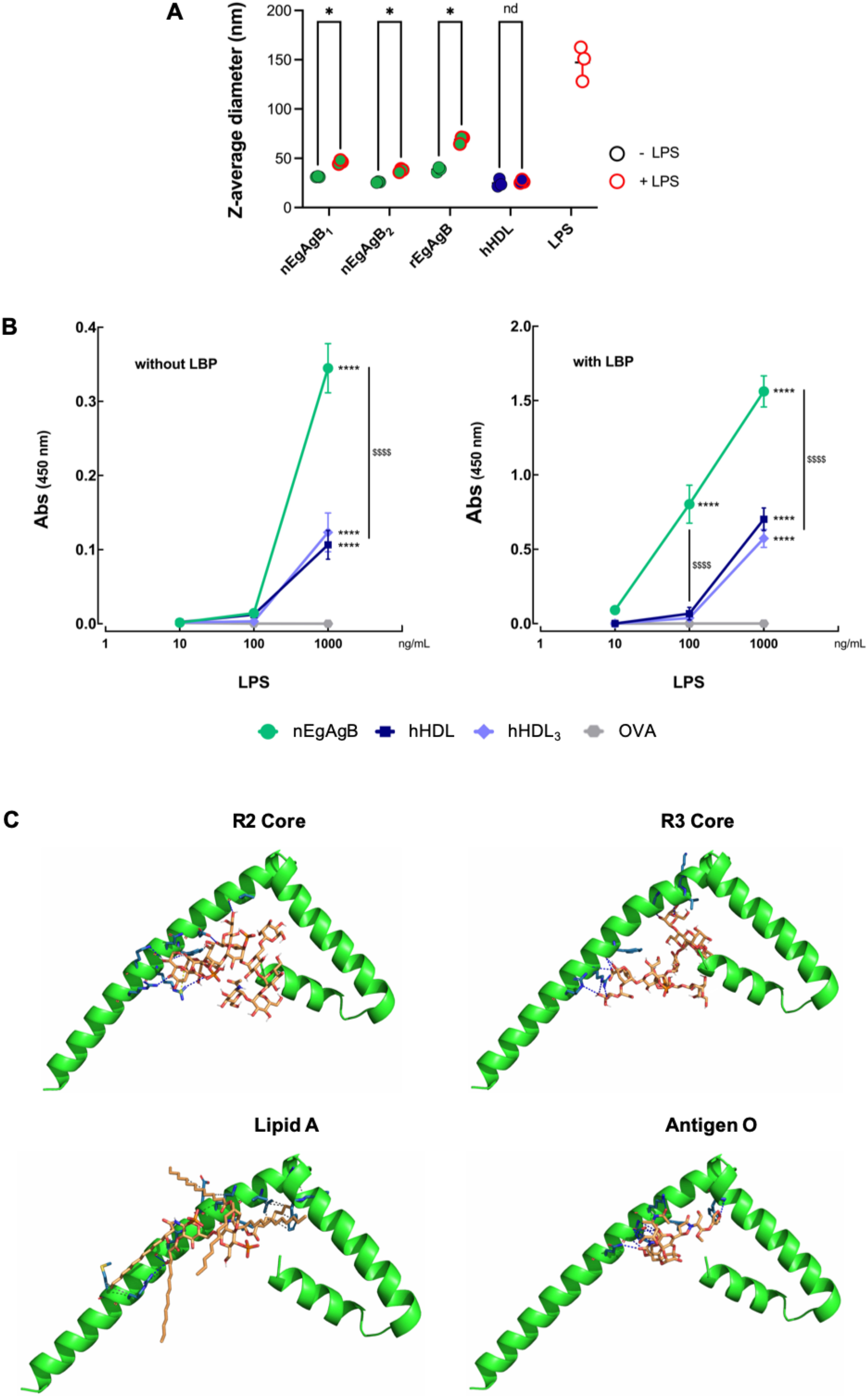
Analysis of the ability of nEgAgB and rEgAgB to bind LPS. A) nEgAgB, rEgAgB or hHDL (all at 1 mg/mL) were mixed with an equal volume of LPS (1 mg/mL) or vehicle (PBS_EB_) and analysed by DLS. In parallel LPS was analysed under the same conditions. Results are shown as the Z-average of three consecutive determinations. Significant differences between the selected conditions are indicated with * (t-test, p< 0.05). B) EgAgB (green circles), total hHDL (hHDL, blue squares), hHDL_3_ (light blue diamond) or OVA (grey hexagon, as irrelevant protein control) were immobilised on ELISA plates at 10 µg/mL each. After blocking with PBS-BSA (1%), biotinylated LPS (1, 10, 100 or 1000 ng/mL) was added in the absence or presence of hLBP (0.1 µg/mL) as indicated. LPS binding was detected with Sv-HRP followed by HRP activity determination. The absorbance at 450 nm (Abs_450_), representing the mean ± SD of data from three independent experiments with analytical duplicates, was plotted against LPS concentration. Significant differences compared to OVA or between the selected conditions are indicated with * and $, respectively (two-way ANOVA and Tukey’s test, ****/$$$$ p < 0.0001). C) Molecular docking analysis using HADOOCK of the interaction between EgAgB8/1 subunit (accession number U6JQF4) and the R2 core and R3 core, O antigen and lipid A regions of LPS.

To confirm an EgAgB interaction with LPS we used a binding assay previously described to assess LPS binding to hHDL and hHDL_3_ (22). hHDL and hHDL_3_ were both capable of binding LPS, assessed in a range of 100 and 1000 ng/mL, with enhanced efficiency when LBP was present (Fig. 9B). EgAgB also bound LPS both in the absence or presence of LBP, demonstrating an increased interaction compared to hHDL/hHDL_3_, as a higher binding was observed at 1000 ng/mL LPS in the absence of LBP, and at 100 and 1000 ng/mL LPS in the presence of LBP. The LPS binding activity of EgAgB was similar for nEgAgB preparations purified with or without ultracentrifugation. Moreover, this activity was found to be thermostable, as it remained unchanged after treatment at 100 °C for 10 min (Supplementary Fig. 7).

In addition, molecular docking analyses were performed using HADDOCK to assess the interaction of EgAgB8/1 with individual structural domains of bacterial LPS, including lipid A, the R2 and R3 core oligosaccharides, and the O-antigen. All LPS regions tested were capable of interacting with EgAgB8/1 (Fig. 9C), forming a substantial number of hydrogen bonds that contributed to the stabilization of the complexes. Among the most recurrent interacting residues were Arg66, Arg70, Glu59, and Phe62, which established multiple hydrogen bonds and salt bridges with the lipid A and R3 core. The O-antigen and R2 core domains exhibited comparatively fewer interactions, with Glu59 and Gln63 as the primary contributors. Hydrophobic contacts with Phe42, Leu47, and Val51 further stabilized the lipid A–EgAgB8/1 complex (Supplementary Table 2). These findings support the existence of a defined LPS-binding interface in EgAgB8/1, with a preferential affinity for the lipid A and core oligosaccharide regions, which may underlie its modulatory effect on LPS-mediated immune activation.

## Discussion

This study provides compelling evidence of a selective binding between EgAgB and LPS, uncovering a novel interaction that likely contributes to the modulatory effects of this parasite lipoprotein on myeloid cells, as shown in previous *in vitro* and *in vivo* studies (11, 12, 14, 17). This was especially apparent given the frequent presence of LPS traces in EgAgB preparations. Consistently, achieving LPS-free nEgAgB has posed a significant challenge throughout our studies. As such, traces of LPS were frequently detected in nEgAgB preparations but not in buffer controls, despite using exigent purification protocols (including an immunoaffinity step based on monoclonal antibodies anti-EgAgB8/1 (14, 39), and stringent measures such as pyrogen-free buffers and the use of laminar flow cabinet for immunoaffinity and desalting steps). The origin of LPS carried by EgAgB remains unclear; it may bind to EgAgB during infection or throughout the purification process. Both scenarios could account for the observed variability in LPS content among nEgAgB batches. LPS derived from the gut microbiota is likely present in infected viscera such as the liver and lungs, although its abundance may vary between hosts depending on several factors, including the metabolic and nutritional status as well as diet (40–42). Additionally, differences in the microbiological quality of the parasite material likely influence the levels of LPS that could be carried from the host viscera into the collected HF. Notably, LPS traces were also detected in rEgAgB expressed in *D. melanogaster* cells, indicating that LPS was retained during purification despite the stringent and controlled conditions used for production and immunopurification. A previous study did not find significant LPS levels in the EgAgB preparation used for studying effects on DC (11); however, this study applied a boiling treatment during purification, which may have inactivated LPS and affected its quantification. Although additional studies on some immunological properties of EgAgB exist, LPS levels in EgAgB preparations purified from HF have not been routinely evaluated (9, 10, 43–45). More recently, some investigations have been performed with EgAgB preparations in which LPS may have been inactivated by heating and/or depleted by adsorption on commercial resins (15, 17), all likely contributing to an underestimated LPS content and undervalue EgAgB’s ability to bind LPS selectively.

The amount of LPS carried by EgAgB might be near the theoretical threshold required to induce inflammatory cytokine secretion in myeloid cells, particularly DC (reviewed in (46)). Data on *in vitro* activation of myeloid cells by exposure to EgAgB as the sole stimulus is limited. In this work nEgAgB and rEgAgB (up to 10 µg/mL) on its own induced secretion of proinflammatory cytokines by BMDC whereas no effects were observed in THP-1 macrophages or BMDM (12, 14, 17). This discrepancy may be explained by the higher sensitivity of BMDC to detect PAMPs compared to macrophages. When analysing whether this activation was attributable to EgAgB-carried LPS, polymyxin B treatment failed to inhibit EgAgB-induced cytokine responses in BMDC suggesting that EgAgB either interferes with polymyxin B’s ability to neutralize LPS or directly engages innate cell signalling pathways. The former explanation is more plausible, as in our hands polymyxin B-Sepharose beads were ineffective at removing LPS from EgAgB due to a high-affinity interaction that hindered EgAgB elution even in the presence of urea. Similar binding properties between human lipoproteins and polymyxin B have been reported (47). Interestingly, BMDC activation by nEgAgB/rEgAgB preparations was TLR4-dependent as a nearly complete reduction in the cytokine responses was observed after pretreatment with the TLR4 inhibitor TAK-242 or using *Tlr4*^-/-^ BMDC. EgAgB itself does not appear to trigger TLR4-mediated signalling in BMDC, since its binding to BMDC occurs independently of TLR4, as demonstrated using WT and *Tlr2^-/-^*and *Tlr4^-/-^* BMDC. Collectively, these results suggest that in our *in vitro* assays EgAgB acted as an LPS carrier, delivering LPS to the MD2/TLR4 complex on BMDC, even at suboptimal or minimally activating doses (Fig. 10A). Furthermore, based on data from *in vitro* binding assays, LBP likely facilitates LPS binding to EgAgB. This might involve LBP extracting LPS monomers from micelles and subsequently transferring them to EgAgB, analogous to its established mechanism with other acceptors, such as hHDL (38, 48).

**Figure 10.**
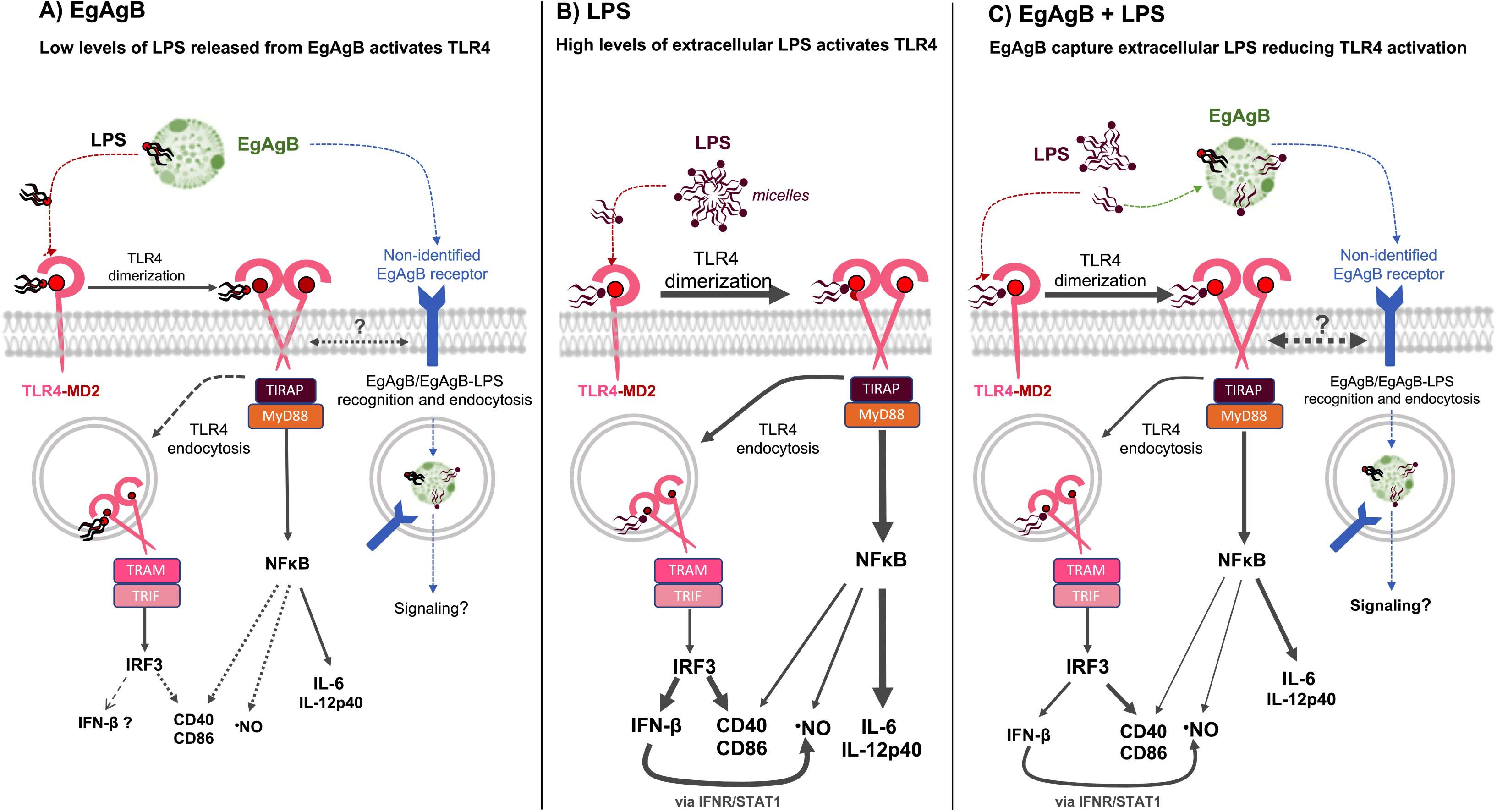
Hypothetical mechanisms involved in EgAgB effects on DC activation. A) In the extracellular environment, EgAgB carries LPS traces, which could be transferred to TLR4/MD2 to trigger a TLR4-mediated inflammatory response in DC. This response would involve TLR4 dimerization and TIRAP/MyD88/NFκB activation, with the consequence production of low levels of IL-6, IL-12p40 and ·NO. TLR4 internalization might lead to a null/limited TRAM/TRIF/IRF3 activation, in agreement with undetectable IFN-ꞵ secretion and slight increases in cell surface costimulatory molecules (CD40, CD86). Putative, non-identified, EgAgB cell receptors may be involved in EgAgB/EgAgB-LPS recognition and additional signalling actions from the plasma membrane or endosomes. B) Scheme of the main intracellular signalling pathways induced in DC by TLR4/MD2 activation by high concentrations of LPS in the extracellular milieu. TLR4/MD2 activation results in TLR4 dimerization and TIRAP/MyD88/NFκB activation, inducing the secretion of high levels of IL-6 and IL-12p40, while TRAM/TRIF/IRF3 activation is responsible for IFN-ꞵ secretion. Both signalling pathways contribute to ·NO and cell surface CD40 and CD86 responses, with an important contribution of IFN-ꞵ to ·NO production, via IFNR/STAT1-mediated autocrine activation. C) In the extracellular environment, EgAgB competes with TLR4/MD2 for LPS binding, capturing LPS and, in turn, interfering with TLR4 dimerization/endocytosis and the consequent cytokine and ·NO responses. In comparison, modulation of CD40 and CD86 would be less efficient, which may be linked to putative EgAgB/EgAgB-LPS recognition and additional signalling actions from the plasma membrane or endosomes.

Whether EgAgB could directly trigger inflammatory signalling pathways in myeloid cells remains to be elucidated. *In vitro* stimulation with EgAgB caused activation of immature (increased surface CD86) and mature (increased surface CD40, IL-6 and TNFɑ secretion) human DC, which was associated with IRAK phosphorylation and NF-κB activation (11). Since in that study, LPS traces were not detected in EgAgB preparations (based on LAL assay) this activation was attributed to a direct EgAgB’s effect. A putative LPS-independent activation of DC by EgAgB aligns with the residual -albeit weak-BMDC activation by nEgAgB observed in our experiments using *Tlr4*^-/-^ BMDC. Lipoprotein receptors could participate in EgAgB-DC interactions, since we previously observed that nEgAgB binding to THP-1 monocytes was partially inhibited by LDL and HDL (12). The structural similarities between EgAgB subunits and LDL/HDL exchangeable apolipoproteins —such as high α-helix content and distribution of polar/charged versus hydrophobic amino acids— may explain why these evolutionarily distant lipoproteins share cell receptors. However, a receptor for EgAgB in myeloid cells has not been identified yet (Fig. 10A); involvement of LDL receptor, LRP1 or SR-A was not supported by competitive binding assays using monocytes (12). Further investigation is warranted to decipher EgAgB targets in DC.

Despite EgAgB preparations on its own induced BMDC activation, they also partially inhibited LPS-induced BMDC cytokine and ·NO responses when co-administered with 10 ng/mL LPS (at least one order of magnitude higher than maximal levels in nEgAgB batches) (Fig. 10C). Interestingly, these responses correlated with a reduction of LPS-induced CD14 expression and interruption of TLR4 dimerization/endocytosis, which are initial steps in the TLR4 activation cascade. In agreement with our results, Zhang *et al.* recently reported that EgAgB inhibits LPS-induced TLR4 dimerization and endocytosis in an artificial cell model using HEK293T cells transfected with CD14, TLR4 and MD2 (17). Considering EgAgB’s ability to bind LPS, our findings suggest that EgAgB modulatory actions, observed here and previously (11–14, 17), may involve extracellular sequestration of LPS. LPS capture in the milieu could also participate in the previously described downregulation effect of heat-treated EgAgB on LPS-induced mature DC activation (11), considering EgAgB binding to LPS resisted boiling for 10 minutes. This complex scenario reveals that EgAgB interaction with LPS might have opposite consequences for myeloid cell activation, at least *in vitro* studies. First, EgAgB may facilitate TLR4/MD2^+^ cell activation by carrying and delivering trace amounts of LPS (Fig. 10A). Conversely, EgAgB may prevent BMDC activation by scavenging high LPS amounts from the extracellular milieu (Fig. 10C). The mentioned opposite outcomes would depend on LPS concentrations and EgAgB-LPS binding affinity. Since LPS comprises a group of structurally similar but non-identical macromolecules (as reviewed by (46)), further studies are needed to determine how LPS biochemical variations influence its binding to EgAgB. On the other hand, as EgAgB belongs to a complex multigenic and polymorphic family (49), its ability to bind and sequester LPS might also differ among EgAgB8 isoforms. EgAgB8/1 is the predominant subunit in nEgAgB purified from HF of various *E. granulosus* s.l. species/genotypes (19), but differences in the subunit composition between parasite stages as well as along the time of the infection in different hosts likely exist (50).

The hypothesis of LPS extracellular sequestration aligns with EgAgB’s modulatory effects on cytokine secretion (IL-6, IL-12p40, IFN-ꞵ) and ·NO generation, observed in this study and previous macrophage research (14, 15, 17). In fact, LPS sequestration would interfere with the activation of all TLR4-mediated signalling pathways triggered by LPS in myeloid cells (Fig. 10B). These include the TIRAP/MyD88 pathway, which leads to early NFκB activation and is primarily responsible for IL-6 and IL-12p40 secretion, and the TRAM/TRIF pathway, which activates IRF3 and results in IFN-ꞵ secretion (51–54). The generation of ·NO requires both TIRAP/MyD88 and TRAM/TRIF pathways, with the latter significantly amplifying *Nos2* upregulation via autocrine IFN-ꞵ (55). Inhibition by EgAgB of the LPS-induced activation of NFκB and IRF3 in macrophages has been recently described (17), in agreement with our interpretation of results. Remarkably, EgAgB exhibited minimal to no modulation of LPS-induced upregulation of cell surface CD40 and CD86, despite these responses also depend on NFκB activation and being enhanced by TRAM/TRIF/IRF3 and IFN-ꞵ/STAT1 in DC (reviewed by (56)). A higher EgAgB’s ability to modulate the inflammatory cytokines compared to costimulatory receptor responses was also reported in LPS-activated macrophages (14). These observations appear contradictory given that CD40 and CD86 upregulation require approximately five-fold higher LPS concentration compared to those needed for inflammatory cytokine secretion (46). Therefore, if EgAgB captures a significant amount of LPS sufficient to inhibit cytokine responses, the residual free LPS levels should be less capable of activating the costimulatory molecule responses, resulting in significant inhibition. The weak or null modulation of cell surface CD40 and CD86 might be due to complementary recognition of EgAgB by cell receptors, which could activate signalling pathways linked to *Cd40/Cd86* expression, thereby counterbalancing modulatory effects from LPS sequestration (Fig. 10C). These putative signalling pathways might involve the recognition and endocytosis of EgAgB/EgAgB-LPS complexes, potentially influencing signalling from the plasma membrane or the endosomal compartment. Endocytosis and subsequent internalization of EgAgB/EgAgB-LPS complexes likely occur given the described EgAgB endocytic uptake by macrophages (37). The modulation of costimulatory molecules in myeloid cells by EgAgB requires further examination, as the presence of an additional stimuli (i.e. IFN-γ) might influence the efficacy of EgAgB to inhibit LPS-driven effects (17).

Considering the similar size of nEgAgB and HDL (14, 18), our results (based on DLS analysis and the ELISA-like binding assay) suggest that the apparent affinity of EgAgB-LPS interaction is higher than that of hHDL/HDL_3_-LPS. This is consistent with the observation that *in vitro* nEgAgB diminished the secretion of inflammatory cytokines at significantly lower concentrations than hHDL, which supports the notion that EgAgB has a higher potential as an LPS scavenger in the extracellular milieu than HDL. Noteworthy, an opposite trend was found when we analysed EgAgB and hHDL effects on LTA-induced BMDC activation, revealing a selectivity for the ligands that these lipoproteins could bind in physiological conditions. Although both LPS and LTA are amphiphilic molecules, differences in their size, density of negative groups and hydrophobic moieties could explain these results. Besides, despite conserved chemical structures, substantial diversity exists among these bacterial components, which may influence the binding capacity of nEgAgB and hHDL/HDL_3_ and consequently affect cell activation. Further studies are required to explore the differential capacity of EgAgB to bind LPS and LTA.

The main protein subunit of nEgAgB, EgAgB8/1, was assembled *in vitro* as a lipoprotein that resembles nEgAgB in all functional assays, although it exhibited slightly lower modulatory activity on LPS-driven BMDC activation. These results were in agreement with our previous study on macrophages (14), supporting rEgAgB8/1 as an interesting model for EgAgB studies despite the impact of differences in lipid composition and putative protein modifications, which might be worth studying. Molecular docking analysis, based on the modelled structure of EgAgB8/1, was consistent with our experimental observations, reinforcing the hypothesis that EgAgB8/1 can directly interact with LPS molecules. Notably, all LPS regions analysed— including lipid A, R2, R3, and the O-antigen—engaged in numerous hydrogen bonds with EgAgB8/1 residues, underscoring the versatility and strength of these interactions. The abundance of hydrogen bonds likely contributes substantially to the affinity and stability of the complexes formed, supporting an interaction that is both specific and structurally meaningful. The prominent involvement of basic and aromatic residues, such as Arg66, Arg70, and Phe62, which are known to mediate lipid and glycan interactions in other apolipoproteins (57, 58), further supports a biologically relevant binding mode. Collectively, these findings provide a structural rationale for the LPS-neutralizing activity of EgAgB1 and its proposed role in modulating host immune response.

The potential of EgAgB to bind and sequester LPS from the extracellular milieu acting as an LPS scavenger macromolecule raises questions about its physiological relevance during CE. LPS has been widely used to model TLR-triggered responses and to study parasite immunomodulation mechanisms in *Echinococcus* (11, 12, 15, 59–61) and other helminthiasis ((62–65) among others) although it is not a helminth-derived PAMP. However, LPS is commonly found in the liver –the most frequent site for hydatid establishment and growth– accounting for 68.8% to 80% of cases (66). Indeed, most luminal LPS can physiologically cross the intestinal barrier, particularly during fat absorption, and subsequently, travel via the portal vein to the liver (67). Although LPS levels in plasma or infected tissues from CE patients have not been evaluated, the contribution of LPS to liver inflammation during infection cannot be ruled out. Furthermore, alterations in the gut microbiota during CE have been observed (68), which might influence both the concentration and pro-inflammatory potential of liver LPS species during infection. Interestingly, EgAgB showed an ability to bind LPS comparable to -if not greater than-hHDL_3_, supporting its putative role as an enteric LPS scavenger in the liver (22). In this scenario, the presence of EgAgB in the vicinity of the hydatid may prevent LPS-mediated inflammation, thus reducing the recruitment of potentially harmful immune cells. Notably, *in vivo* administration of EgAgB was found to protect against pathology in murine models of colitis and sepsis by caecal ligation and puncture (CLP), likely involving modulation of the differentiation of macrophage populations towards less inflammatory, M2-like phenotypes (15, 16). In both models, systemic increases in LPS levels are expected. Our findings suggest that EgAgB protective effects in these models may include LPS sequestration, limiting TLR4-dependent activation pathways in immune cells and, thereby, inhibiting macrophage differentiation into classic pro-inflammatory phenotypes.

In sum, this work sheds light on a novel binding property of EgAgB that may represent a potential parasite strategy to mitigate host inflammation against the hydatid. The multigenic and polymorphic nature of the EgAgB family could broaden the spectrum of LPS species that can be recognized and bound. Whether EgAgB interacts with other PAMPs remains to be fully investigated; however, its failure to inhibit LTA-trigger BMDC activation suggests a more selective recognition profile than HDL. It is worth noting that EgAgB’s ability to bind LPS does not account for other modulation effects on myeloid cells, such as the inhibition of the activation and/or chemotaxis of neutrophils (9, 10, 44) and monocytes (4), and the FcR-mediated phagocytosis by macrophages (17). Thus, further studies are warranted to elucidate, at the molecular level, additional EgAgB mechanisms with regulatory effects on myeloid cells.

## Acknowledgements

The authors thank to M.Sc. Carlos González (Montevideo, Uruguay) for assistance with the statistical analysis.

## References

1. Breijo M, Anesetti G, Martínez L, Sim RB, Ferreira AM. 2008. Echinococcus granulosus: the establishment of the metacestode is associated with control of complement-mediated early inflammation. 2. Exp Parasitol 118:188–196.

2. Díaz A, Casaravilla C, Barrios AA, Ferreira AM. 2016. Parasite molecules and host responses in cystic echinococcosis. 3. Parasite Immunology 38:193–205.

3. Siracusano A, Delunardo F, Teggi A, Ortona E. 2012. Host-parasite relationship in cystic echinococcosis: an evolving story. Clinical & developmental immunology 2012:639362.

4. Silva-Álvarez V, Maite A, Lía A, Zamarreño F, Costabel D, García-zepeda E, Salinas G, Córsico B, María A. 2015. Echinococcus granulosus antigen B : A Hydrophobic Ligand Binding Protein at the host – parasite interface. Prostaglandins Leukotrienes and Essential Fatty Acids 93:17–23.

5. Kronenberg PA, Reinehr M, Eichenberger RM, Hasler S, Laurimäe T, Weber A, Deibel A, Müllhaupt B, Gottstein B, Müller N, Hemphill A, Deplazes P. 2023. Monoclonal antibody-based localization of major diagnostic antigens in metacestode tissue, excretory/secretory products, and extracellular vesicles of Echinococcus species. Front Cell Infect Microbiol 13:1162530.

6. Lorenzo C, Ferreira HB, Monteiro KM, Rosenzvit M, Kamenetzky L, García HH, Vasquez Y, Naquira C, Sánchez E, Lorca M, Contreras M, Last JA, González-Sapienza GG. 2005. Comparative analysis of the diagnostic performance of six major Echinococcus granulosus antigens assessed in a double-blind, randomized multicenter study. 6. Journal of clinical microbiology 43:2764–2770.

7. Ortona E, Riganò R, Margutti P, Notargiacomo S, Ioppolo S, Vaccari S, Barca S, Buttari B, Profumo E, Teggi A, Siracusano A. 2000. Native and recombinant antigens in the immunodiagnosis of human cystic echinococcosis. 11. Parasite immunology 22:553–9.

8. Carmena D, Benito A, Eraso E. 2006. Antigens for the immunodiagnosis of Echinococcus granulosus infection: An update. 1. Acta tropica 98:74–86.

9. Shepherd JC, Aitken A, McManus DP. 1991. A protein secreted in vivo by Echinococcus granulosus inhibits elastase activity and neutrophil chemotaxis. 1. Molecular and biochemical parasitology 44:81–90.

10. Riganò R, Profumo E, Bruschi F, Carulli G, Azzarà A, Ioppolo S, Buttari B, Ortona E, Margutti P, Teggi A, Siracusano A, Azzara A. 2001. Modulation of Human Immune Response by Echinococcus granulosus Antigen B and Its Possible Role in Evading Host Defenses. 1. Infection and immunity 69:288–296.

11. Riganò R, Buttari B, Profumo E, Ortona E, Delunardo F, Margutti P, Mattei V, Teggi A, Sorice M, Siracusano A. 2007. Echinococcus granulosus antigen B impairs human dendritic cell differentiation and polarizes immature dendritic cell maturation towards a Th2 cell response. 4. Infection and immunity 75:1667–78.

12. Silva-Álvarez V, Folle AM, Ramos ALL, Kitano ES, Iwai LK, Corraliza I, Córsico B, Ferreira AMM. 2016. Echinococcus granulosus Antigen B binds to monocytes and macrophages modulating cell response to inflammation. 1. Parasites & vectors 9:69.

13. Silva-Álvarez V, Ramos AL, Folle AM, Lagos S, Dee VM, Ferreira AM. 2018. Antigen B from Echinococcus granulosus is a novel ligand for C-reactive protein. 9. Parasite Immunology 40:e12575.

14. Folle AM, Lagos Magallanes S, Fló M, Alvez-Rosado R, Carrión F, Vallejo C, Watson D, Julve J, González-Sapienza G, Pristch O, González-Techera A, Ferreira AM. 2024. Modulatory actions of Echinococcus granulosus antigen B on macrophage inflammatory activation. Front Cell Infect Microbiol 14:1362765.

15. Bao J, Qi W, Sun C, Tian M, Jiao H, Guo G, Guo B, Ren Y, Zheng H, Wang Y, Yan M, Zhang Z, McManus DP, Li J, Zhang W. 2022. Echinococcus granulosus sensu stricto and antigen B may decrease inflammatory bowel disease through regulation of M1/2 polarization. Parasites & Vectors 15:391.

16. Qian Y-Y, Huang F-F, Chen S-Y, Zhang W-X, Wang Y, Du P-F, Li G, Ding W-B, Qian L, Zhan B, Chu L, Jiang D-H, Yang X-D, Zhou R. 2024. Therapeutic effect of recombinant Echinococcus granulosus antigen B subunit 2 protein on sepsis in a mouse model. Parasites Vectors 17:467.

17. Zhang Y, Yue Y, Cheng Y, Jiao H, Yan M. 2025. Antigen B from Echinococcus granulosus regulates macrophage phagocytosis by controlling TLR4 endocytosis in immune thrombocytopenia. Chem Biol Interact 406:111350.

18. Obal G, Ramos AL, Silva V, Lima A, Batthyany C, Bessio MI, Ferreira F, Salinas G, Ferreira AM. 2012. Characterisation of the native lipid moiety of Echinococcus granulosus antigen B. 5. PLoS neglected tropical diseases 6:e1642.

19. Folle AM, Kitano ES, Lima A, Gil M, Cucher M, Mourglia-Ettlin G, Iwai LK, Rosenzvit M, Batthyány C, Ferreira AM. 2017. Characterisation of Antigen B Protein Species Present in the Hydatid Cyst Fluid of Echinococcus canadensis G7 Genotype. 1. PLoS Neglected Tropical Diseases 11:e0005250.

20. Chapman MJ. 1980. Animal lipoproteins: chemistry, structure, and comparative aspects. 7. Journal of lipid research 21:789–853.

21. Meilhac O, Tanaka S, Couret D. 2020. High-Density Lipoproteins Are Bug Scavengers. Biomolecules 10:598.

22. Han Y-H, Onufer EJ, Huang L-H, Sprung RW, Davidson WS, Czepielewski RS, Wohltmann M, Sorci-Thomas MG, Warner BW, Randolph GJ. 2021. Enterically derived high-density lipoprotein restrains liver injury through the portal vein. Science 373:eabe6729.

23. Cucher M, Prada L, Mourglia-Ettlin G, Dematteis S, Camicia F, Asurmendi S, Rosenzvit M. 2011. Identification of Echinococcus granulosus microRNAs and their expression in different life cycle stages and parasite genotypes. 3. International Journal for Parasitology 41:439–.

24. Lutz MB, Kukutsch N, Ogilvie AL, Rössner S, Koch F, Romani N, Schuler G. 1999. An advanced culture method for generating large quantities of highly pure dendritic cells from mouse bone marrow. J Immunol Methods 223:77–92.

25. Havel RJ, Eder HA, Bragdon JH. 1955. The distribution and chemical composition of ultracentrifugally separated lipoproteins in human serum. J Clin Invest 34:1345–1353.

26. Grisham MB, Johnson GG, Lancaster JR. 1996. Quantitation of nitrate and nitrite in extracellular fluids. Methods Enzymol 268:237–246.

27. Tan Y, Zanoni I, Cullen TW, Goodman AL, Kagan JC. 2015. Mechanisms of Toll-like receptor 4 endocytosis reveal a common immune-evasion strategy used by pathogenic and commensal bacteria. Immunity 43:909–922.

28. Zoete V, Cuendet MA, Grosdidier A, Michielin O. 2011. SwissParam: a fast force field generation tool for small organic molecules. J Comput Chem 32:2359–2368.

29. O’Boyle NM, Banck M, James CA, Morley C, Vandermeersch T, Hutchison GR. 2011. Open Babel: An open chemical toolbox. J Cheminform 3:33.

30. Jumper J, Evans R, Pritzel A, Green T, Figurnov M, Ronneberger O, Tunyasuvunakool K, Bates R, Žídek A, Potapenko A, Bridgland A, Meyer C, Kohl SAA, Ballard AJ, Cowie A, Romera-Paredes B, Nikolov S, Jain R, Adler J, Back T, Petersen S, Reiman D, Clancy E, Zielinski M, Steinegger M, Pacholska M, Berghammer T, Bodenstein S, Silver D, Vinyals O, Senior AW, Kavukcuoglu K, Kohli P, Hassabis D. 2021. Highly accurate protein structure prediction with AlphaFold. Nature 596:583–589.

31. van Zundert GCP, Rodrigues JPGLM, Trellet M, Schmitz C, Kastritis PL, Karaca E, Melquiond ASJ, van Dijk M, de Vries SJ, Bonvin AMJJ. 2016. The HADDOCK2.2 Web Server: User-Friendly Integrative Modeling of Biomolecular Complexes. J Mol Biol 428:720–725.

32. Salentin S, Schreiber S, Haupt VJ, Adasme MF, Schroeder M. 2015. PLIP: fully automated protein-ligand interaction profiler. Nucleic Acids Res 43:W443–447.

33. Domingues MM, Inácio RG, Raimundo JM, Martins M, Castanho MARB, Santos NC. 2012. Biophysical characterization of polymyxin B interaction with LPS aggregates and membrane model systems. Biopolymers 98:338–344.

34. Puig N, Montolio L, Camps-Renom P, Navarra L, Jiménez-Altayó F, Jiménez-Xarrié E, Sánchez-Quesada JL, Benitez S. 2020. Electronegative LDL Promotes Inflammation and Triglyceride Accumulation in Macrophages. Cells 9:583.

35. Bagheri B, Khatibiyan Feyzabadi Z, Nouri A, Azadfallah A, Mahdizade Ari M, Hemmati M, Darban M, Alavi Toosi P, Banihashemian SZ. 2024. Atherosclerosis and Toll-Like Receptor4 (TLR4), Lectin-Like Oxidized Low-Density Lipoprotein-1 (LOX-1), and Proprotein Convertase Subtilisin/Kexin Type9 (PCSK9). Mediators Inflamm 2024:5830491.

36. Korbecki J, Bajdak-Rusinek K. 2019. The effect of palmitic acid on inflammatory response in macrophages: an overview of molecular mechanisms. Inflamm Res 68:915–932.

37. Danieli Da Silva E, Cancela M, Monteiro KM, Ferreira HB, Zaha A, da Silva ED, Cancela M, Monteiro KM, Ferreira HB, Zaha A. 2018. Antigen B from Echinococcus granulosus enters mammalian cells by endocytic pathways. 5. PLoS Neglected Tropical Diseases 12:e0006473.

38. Ryu J-K, Kim SJ, Rah S-H, Kang JI, Jung HE, Lee D, Lee HK, Lee J-O, Park BS, Yoon T-Y, Kim HM. 2017. Reconstruction of LPS Transfer Cascade Reveals Structural Determinants within LBP, CD14, and TLR4-MD2 for Efficient LPS Recognition and Transfer. Immunity 46:38–50.

39. González G, Nieto A, Fernández C, Orn A, Wernstedt C, Hellman U. 1996. Two different 8 kDa monomers are involved in the oligomeric organization of the native Echinococcus granulosus antigen B. 12. Parasite immunology 18:587–96.

40. Tang J, Xu L, Zeng Y, Gong F. 2021. Effect of gut microbiota on LPS-induced acute lung injury by regulating the TLR4/NF-kB signaling pathway. Int Immunopharmacol 91:107272.

41. Yang J, Liu J, Gu H, Song W, Zhang H, Wang J, Yang P. 2025. Gut microbiota, metabolites, and pulmonary hypertension: Mutual regulation and potential therapies. Microbiological Research 299:128245.

42. Icaza-Chávez ME. 2013. Microbiota intestinal en la salud y la enfermedad. Revista de Gastroenterología de México 78:240–248.

43. Riganò R, Buttari B, De Falco E, Profumo E, Ortona E, Margutti P, Scottà C, Teggi A, Siracusano A. 2004. Echinococcus granulosus-specific T-cell lines derived from patients at various clinical stages of cystic echinococcosis. 1. Parasite Immunology 26:45–52.

44. Virginio VG, Taroco L, Ramos AL, Ferreira AM, Zaha A, Ferreira HB, Hernández A. 2007. Effects of protoscoleces and AgB from Echinococcus granulosus on human neutrophils: Possible implications on the parasite’s immune evasion mechanisms. 5. Parasitology Research 100:935–942.

45. Petrone L, Vanini V, Petruccioli E, Ettorre GM, Busi Rizzi E, Schininà V, Girardi E, Ludovisi A, Gómez-Morales MÁ, Pozio E, Teggi A, Goletti D. 2015. IL-4 specific-response in whole blood associates with human Cystic Echinococcosis and cyst activity. Journal of Infection 70:299–306.

46. Bonhomme D, Cavaillon J-M, Werts C. 2023. The dangerous liaisons in innate immunity involving recombinant proteins and endotoxins: Examples from the literature and the Leptospira field. J Biol Chem 300:105506.

47. Liao W, Florén CH. 1993. Polymyxin B complexes with and cationizes low density lipoproteins. The cause of polymyxin B-induced enhancement of endocytotic catabolism of low density lipoproteins. Biochem Pharmacol 45:1835–1843.

48. Wurfel MM, Kunitake ST, Lichenstein H, Kane JP, Wright SD. 1994. Lipopolysaccharide (LPS)-binding protein is carried on lipoproteins and acts as a cofactor in the neutralization of LPS. J Exp Med 180:1025–1035.

49. Olson PD, Zarowiecki M, Kiss F, Brehm K. 2012. Cestode genomics -progress and prospects for advancing basic and applied aspects of flatworm biology. 2–3. Parasite immunology 34:130–50.

50. Zhang W, Li J, Jones MK, Zhang Z, Zhao L, Blair D, McManus DP. 2010. The Echinococcus granulosus antigen B gene family comprises at least 10 unique genes in five subclasses which are differentially expressed. 8. PLoS neglected tropical diseases 4:e784.

51. Fitzgerald KA, Rowe DC, Barnes BJ, Caffrey DR, Visintin A, Latz E, Monks B, Pitha PM, Golenbock DT. 2003. LPS-TLR4 signaling to IRF-3/7 and NF-kappaB involves the toll adapters TRAM and TRIF. J Exp Med 198:1043–1055.

52. Kagan JC, Medzhitov R. 2006. Phosphoinositide-Mediated Adaptor Recruitment Controls Toll-like Receptor Signaling. Cell 125:943–955.

53. Kagan JC, Su T, Horng T, Chow A, Akira S, Medzhitov R. 2008. TRAM couples endocytosis of Toll-like receptor 4 to the induction of interferon-β. Nat Immunol 9:361– 368.

54. Tsukamoto H, Takeuchi S, Kubota K, Kobayashi Y, Kozakai S, Ukai I, Shichiku A, Okubo M, Numasaki M, Kanemitsu Y, Matsumoto Y, Nochi T, Watanabe K, Aso H, Tomioka Y. 2018. Lipopolysaccharide (LPS)-binding protein stimulates CD14-dependent Toll-like receptor 4 internalization and LPS-induced TBK1-IKKɛ-IRF3 axis activation. J Biol Chem 293:10186–10201.

55. Fujihara M, Ito N, Pace JL, Watanabe Y, Russell SW, Suzuki T. 1994. Role of endogenous interferon-beta in lipopolysaccharide-triggered activation of the inducible nitric-oxide synthase gene in a mouse macrophage cell line, J774. Journal of Biological Chemistry 269:12773–12778.

56. Ciesielska A, Matyjek M, Kwiatkowska K. 2021. TLR4 and CD14 trafficking and its influence on LPS-induced pro-inflammatory signaling. Cell Mol Life Sci 78:1233–1261.

57. Meyers NL, Larsson M, Vorrsjö E, Olivecrona G, Small DM. 2017. Aromatic residues in the C terminus of apolipoprotein C-III mediate lipid binding and LPL inhibition. J Lipid Res 58:840–852.

58. Anbarasu A, Sethumadhavan R. 2007. Exploring the role of cation-pi interactions in glycoproteins lipid-binding proteins and RNA-binding proteins. J Theor Biol 247:346–353.

59. Casaravilla C, Pittini A, Rückerl D, Seoane PI, Jenkins SJ, MacDonald AS, Ferreira AM, Allen JE, Díaz A. 2014. Unconventional maturation of dendritic cells induced by particles from the laminated layer of larval Echinococcus granulosus. 8. Infection and immunity 82:3164–76.

60. Pittini Á, Martínez-Acosta YE, Casaravilla C, Seoane PI, Rückerl D, Quijano C, Allen JE, Díaz Á. 2019. Particles from the Echinococcus granulosus Laminated Layer Inhibit CD40 Upregulation in Dendritic Cells by Interfering with Akt Activation. Infect Immun 87:e00641–19.

61. Casaravilla C, Pittini Á, Rückerl D, Allen JE, Díaz Á. 2020. Activation of the NLRP3 Inflammasome by Particles from the Echinococcus granulosus Laminated Layer. Infect Immun 88:e00190–20.

62. Harnett W, Harnett MM, Byron O. 2003. Structural/functional aspects of ES-62--a secreted immunomodulatory phosphorylcholine-containing filarial nematode glycoprotein. Curr Protein Pept Sci 4:59–71.

63. Balic A, Smith KA, Harcus Y, Maizels RM. 2009. Dynamics of CD11c+ dendritic cell subsets in lymph nodes draining the site of intestinal nematode infection. Immunol Lett 127:68–75.

64. Terrazas CA, Alcántara-Hernández M, Bonifaz L, Terrazas LI, Satoskar AR. 2013. Helminth-excreted/secreted products are recognized by multiple receptors on DCs to block the TLR response and bias Th2 polarization in a cRAF dependent pathway. FASEB J 27:4547–4560.

65. Hotez PJ, Brindley PJ, Bethony JM, King CH, Pearce EJ, Jacobson J. 2008. Helminth infections: the great neglected tropical diseases. 4. The Journal of clinical investigation 118:1311–21.

66. Salamone G, Licari L, Randisi B, Falco N, Tutino R, Vaglica A, Gullo R, Porrello C, Cocorullo G, Gulotta G. 2016. Uncommon localizations of hydatid cyst. Review of the literature. G Chir 37:180–185.

67. Akiba Y, Maruta K, Takajo T, Narimatsu K, Said H, Kato I, Kuwahara A, Kaunitz JD. 2020. Lipopolysaccharides transport during fat absorption in rodent small intestine. Am J Physiol Gastrointest Liver Physiol 318:G1070–G1087.

68. Cao D, Pang M, Wu D, Chen G, Peng X, Xu K, Fan H. 2022. Alterations in the Gut Microbiota of Tibetan Patients With Echinococcosis. Front Microbiol 13:860909.

